# Reconstructing the invasion route of DNA transposons using extant population samples

**DOI:** 10.1101/729889

**Authors:** Lukas Weilguny, Christos Vlachos, Divya Selvaraju, Robert Kofler

## Abstract

Reconstructing invasion routes of transposable elements (TEs), so far, required capturing an ongoing invasion with population samples from different geographic regions and time points. Here, we propose a more accessible approach. Abundantly occurring internal deletions of DNA transposons allow to trace the direction as well as the path of an invasion, even hundreds of generations after the spread of a TE. We validated this hypothesis with computer simulations and by accurately reproducing the route of the P-element invasion in *Drosophila melanogaster*. Finally, we used our method to shed light on the controversial hobo invasion in *D. melanogaster*. Our approach solely requires sequenced samples from extant populations and sequences of TEs of interest. Hence, DNA transposons in a wide range of model and non-model organisms may be analyzed. Our approach will further our understanding of TE dynamics, migration patterns, and the ecology of species.

## Introduction

Transposable elements (TEs) are short stretches of DNA that selfishly proliferate within host genomes. TE insertions may be grouped into families by the degree of sequence similarity [Wicker et al., 2007]. Two classes of TE families are usually distinguished: families that have a RNA-intermediate (RNA transposons) and families that don’t (DNA transposons) [Finnegan, 1989, Wicker et al., 2007]. DNA transposons mostly propagate by a cut-and-paste mechanism while RNA transposons rely on a copy-and-paste mechanism [Finnegan, 1989, Wicker et al., 2007]. The cut-and-paste replication strategy of DNA transposons does not inherently lead to an increase copy numbers. The copy number increase is achieved by repair of double stranded breaks - resulting from excision of the TE - using the sister chromatid as template [Engels et al., 1990, Daniels and Chovnick, 1993, Gloor et al., 1991]. Interruption of this gap-repair mechanism leads to insertions with internal deletions (IDs) [Engels et al., 1990, Daniels and Chovnick, 1993, Gloor et al., 1991]. These ID elements are usually non-autonomous and require the enzymes encoded by autonomous full-length (FL) insertions for mobilization [Hua-Van et al., 2011, Wicker et al., 2007].

Most TE insertions likely have negative effects on host-fitness [Mackay, 1989, Houle and Nuzhdin, 2004, Mackay et al., 1991, Blumenstiel et al., 2014, Yukuhiro et al., 1985]. To control the spread of these deleterious elements host organism evolved elaborate defence mechanism. In mammals and many invertebrates the defence system against TEs relies on the so called piRNAs, small RNAs ranging in size from 23 to 29nt [Brennecke et al., 2007, Gunawardane et al., 2007]. piRNAs associate with PIWI clade proteins, which act to silence TEs with complementary sequences at the transcriptional as well as the post-transcriptional level [Sienski et al., 2012, Le Thomas et al., 2013, Brennecke et al., 2007, Gunawardane et al., 2007]. piRNAs are largely derived from discrete genomic loci that have been termed piRNA clusters [Brennecke et al., 2007, Malone et al., 2009]. In spite of these elaborate defence mechanism TEs are successful invaders that have been found in most prokaryotic and eukaryotic genomes studied so far [Biémont and Vieira, 2006, Wicker et al., 2007]. The broad taxonomic distribution of TEs is partly due frequent horizontal transfer (HT) of TEs among species, which may trigger a TE invasion in a naive, hitherto uninfected species [Schaack et al., 2010, Peccoud et al., 2017]. For example the P-element, a widely studied DNA transposon, invaded two different Drosophila species within the last 100 years [Kidwell, 1983, Kofler et al., 2015]. Following HT, a novel TE may multiply within populations until the spread is stopped by TE copies that randomly jumped into piRNA clusters, which triggers the production of piRNAs that silence the TE [Bergman et al., 2006, Malone and Hannon, 2009, Zanni et al., 2013, Yamanaka et al., 2014, Goriaux et al., 2014, Duc et al., 2019]. Initially the TE will be stopped by multiple segregating piRNA cluster insertions but at later stages fixed cluster insertions are expected to emerge [Kofler, 2019]. TEs may not only spread within but also between populations. Starting from the origin of the HT event, TEs may gradually infect neighboring populations until eventually worldwide populations acquired the TE. A classic example is the spread of the P-element in *Drosophila melanogaster* populations [Anxolabéhère et al., 1985, Anxolabéhère et al., 1988]. *D. melanogaster* likely acquired the P-element by HT from the distantly related *D. willistoni* in South America [Daniels and Chovnick, 1993]. The P-element first spread in American populations and later invaded European and African populations [Anxolabéhère et al., 1985, Anxolabéhère et al., 1988].

Reconstructing the invasion route of the P-element, however, required two serendipitous events: a very recent invasion of the P-element in *D. melanogaster* and the availability of many fly strains sampled over the course of the P-element invasion from different geographic locations [Anxolabéhère et al., 1985, Anxolabéhère et al., 1988]. However, such time series data are only available for very few species, and even fewer of these data will capture a TE invasion. Alternatively, repeated sampling of populations could be specifically initiated for the purpose of capturing an ongoing TE invasion. Unfortunately, TE invasions are rare and mostly not discovered until a substantial fraction of worldwide populations have been infected, i.e. when it is already too late to commence sampling of time series data [e.g. Kofler et al., 2015, Pascua and Periquet, 1991, Jordan et al., 1999, Bucheton et al., 1992]. Due to these substantial difficulties, we solely know the invasion routes of very few TEs. A simple approach for reconstructing TE invasions would thus be of enormous benefit. Here, we propose that it may be feasible to reconstruct the invasion route of DNA transposons using IDs of TEs as marker. IDs have some properties that make them highly attractive for this task. First, IDs emerge at a high rate. For example, during an experimental P-element invasion at least 140 different IDs emerged within 60 generations [Kofler et al., 2018]. Second, the breakpoints of IDs are mostly random [Kofler et al., 2018]. It is thus unlikely that an identical pair of breakpoints emerges multiple times independently. Third, IDs solely emerge when a TE is active [Engels, 1989], since IDs of DNA transposons result from interruption of gap-repair following transposition of the TE [Engels et al., 1990, Daniels and Chovnick, 1993, Gloor et al., 1991]. We show that each independently invaded population receives a unique set of IDs, i.e. an ID fingerprint, and that this ID fingerprint persists for some time. Based on the two insights, that the fraction of IDs increases in successively invaded populations and that successively invaded populations share similar ID fingerprints, the invasion route of DNA transposons can be reconstructed. We support our approach with a simple model and show that invasion routes can be traced hundreds of generations following the spread of a TE. Additionally, we validated our approach by reconstructing the well known invasion of the P-element in *D. melanogaster*. Finally, we shed light on the controversial invasion route of hobo in *D. melanogaster*. Since solely sequencing data from extant populations and sequences of TEs of interest are required, our approach may be used to trace invasion routes of different DNA transposons in model as well as non-model organisms.

## Results

Here, we propose that the spatial invasion history of DNA transposons, i.e. the invasion route, may be reconstructed using IDs of the transposon as marker. We initially explore this idea with the P-element in Drosophila because i) the P-element has been extensively studied, therefore the genetics and biochemistry of the P-element are well understood [Kelleher, 2016, Majumdar and Rio, 2015, Burt and Trivers, 2008] ii) sequencing data of flies collected from different geographic origins are publicly available [Kapun et al., 2018, Machado et al., 2018, Bergland et al., 2014, Lack et al., 2015] and iii) the invasion route of the P-element in natural populations of *D. melanogaster* is roughly known [Anxolabéhère et al., 1985, Anxolabéhère et al., 1988], which permits validating our approach. Our approach requires estimates of the abundance of IDs for TE families and samples (usually populations) of interest. For this purpose we previously developed a novel tool, DeviaTE [Weilguny and Kofler, 2019]. Briefly, DeviaTE aligns reads of a sample to consensus sequences of TEs (e.g. the P-element) using a local alignment algorithm, reconstructs the breakpoints of IDs from the partial alignment of reads, and finally infers the frequency of IDs by relating the number of reads supporting an ID to the total coverage of the TE (Fig. 1A; [Weilguny and Kofler, 2019]). This approach is indifferent to the genomic insertion sites of TEs. Therefore a reference assembly is not required. We refer to the set of breakpoints of IDs in a sample, together with the frequencies of these IDs as the “ID fingerprint”. As an example, the ID fingerprint of the P-element in a natural population from North America is shown in Fig. 1B.

**Figure 1:**
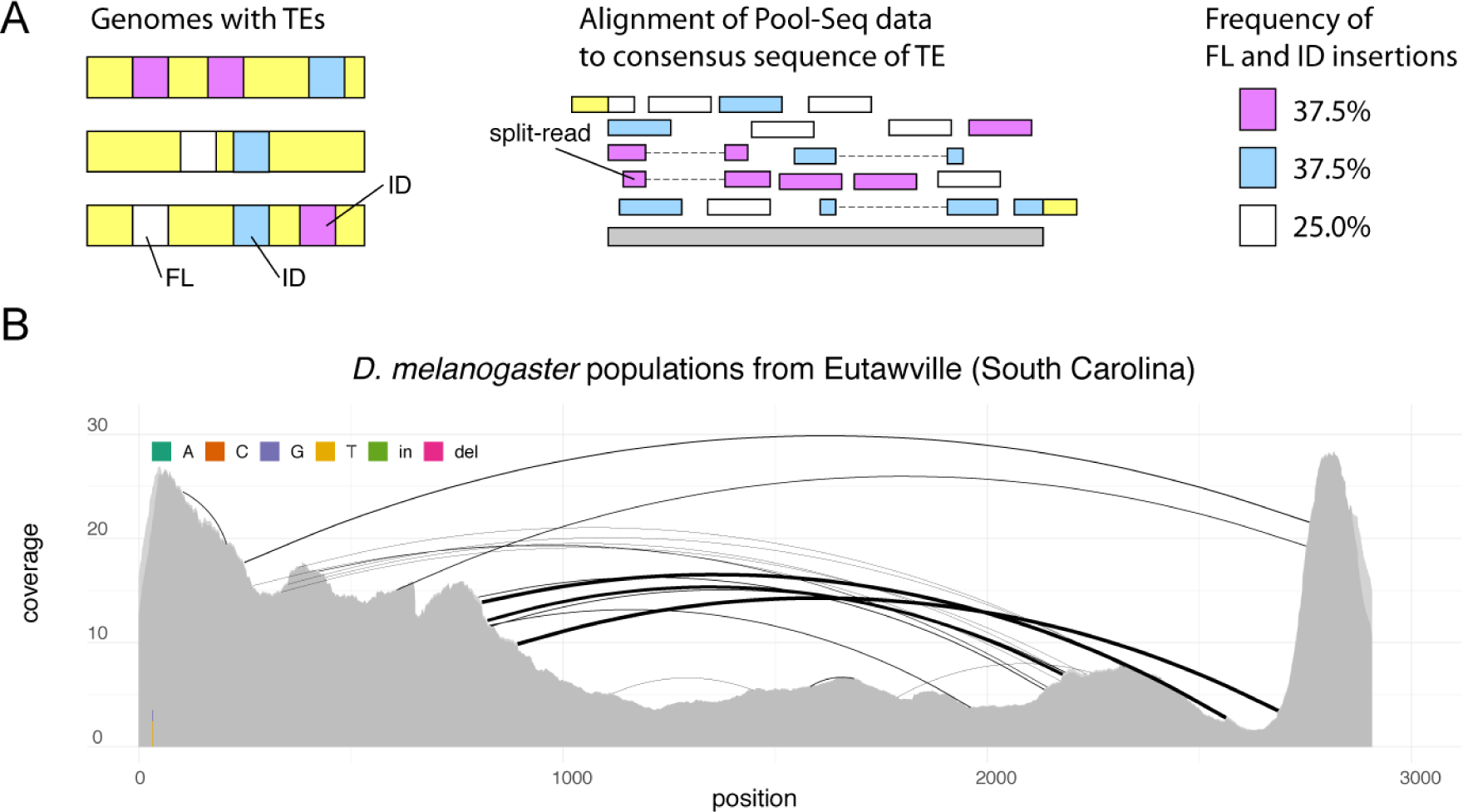
Overview of our approach for taking the ID (internal deletion) fingerprint of a TE family. A) A population harboring some FL (full-length, white) and some ID elements with different breakpoints (blue, magenta; left panel) is sequenced as pool. The reads are aligned to the consensus sequence of a TE family (middle panel). IDs lead to split reads. The frequency of ID and FL insertions can be estimated from the coverage of the TE and the abundance of the split-reads, e.g. using our novel tool DeviaTE (right panel) [Weilguny and Kofler, 2019]. B) Example of an ID fingerprint for the P-element in a *D. melanogaster* population collected 2018 from Eutawville [data from Bergland et al., 2014]. The arcs indicate the breakpoints of IDs and the width of the arcs scales with the frequency of an ID. Three highly abundant IDs (bold arcs) and several less abundant IDs were found.

Ideally, a marker for reconstructing the invasion route of a TE should be characteristic to each invaded population and, in the absence of migration, persist within populations. This requires a marker with two seemingly contradictory properties: i) a high mutation rate during an invasion and ii) a low mutation rate thereafter. IDs of DNA transposons, such as the P-element, may have exactly these two properties since IDs are generated when repair of double-stranded gaps, resulting from excision of a TE, is interrupted [Engels et al., 1990, Daniels and Chovnick, 1993]. The emergence of IDs thus requires TE activity [Engels, 1989, Kofler et al., 2018]. To test if IDs have the two desired properties, we utilized publicly available data of a P-element invasion in experimentally evolving *D. simulans* populations [Kofler et al., 2018]. The authors monitored the spread of the P-element in three replicates by sequencing the populations at each 10^*th*^ generation as pools [Kofler et al., 2018]. P-element copy numbers rapidly increased for the first 20 generations, whereas no further increase was observed during the next 40 generations (Fig. 2A). We first tested whether IDs that emerged during the invasions are characteristic to each replicate. We solely considered IDs that were supported by at least two reads and allowed for a tolerance of 3 bp in the estimated position of ID breakpoints, as exact alignments with indels are frequently not feasible (the position of the gap may be ambiguous). The vast majority of IDs were indeed unique to each replicate (Fig. 2B). Only two out of 47 IDs were found in multiple replicates (Fig. 2B). These two IDs did not necessarily emerge independently in several replicates as they may have already been present in the base population [Kofler et al., 2018]. Next, we tested if IDs persist within populations. Randomly picked IDs were considerably more often recovered at different time points of the same replicate than at any time point of different replicates (Fig. 2C). Finally, we reasoned that by treating each ID as an allele of the TE, it may be feasible to calculate a genetic distance among populations, that reflects the invasion histories of the populations. We used Jost’s D [Jost, 2008] to estimate genetic distances among samples, and the bionj algorithm to construct a tree from the resulting distance matrix [Gascuel, 1997]. Except for the early stage of the invasions, where only very few IDs were found (generation ≤10), all samples were assigned to replicate-specific clades (Fig. 2D). A permutation test, which randomly distributed the 15 samples (≥20 generations) among three different clades (5 samples for each clade) showed that this replicate-specific clustering is highly significant (10^8^ permutations; *p* = 7.8*e* − 06).

**Figure 2:**
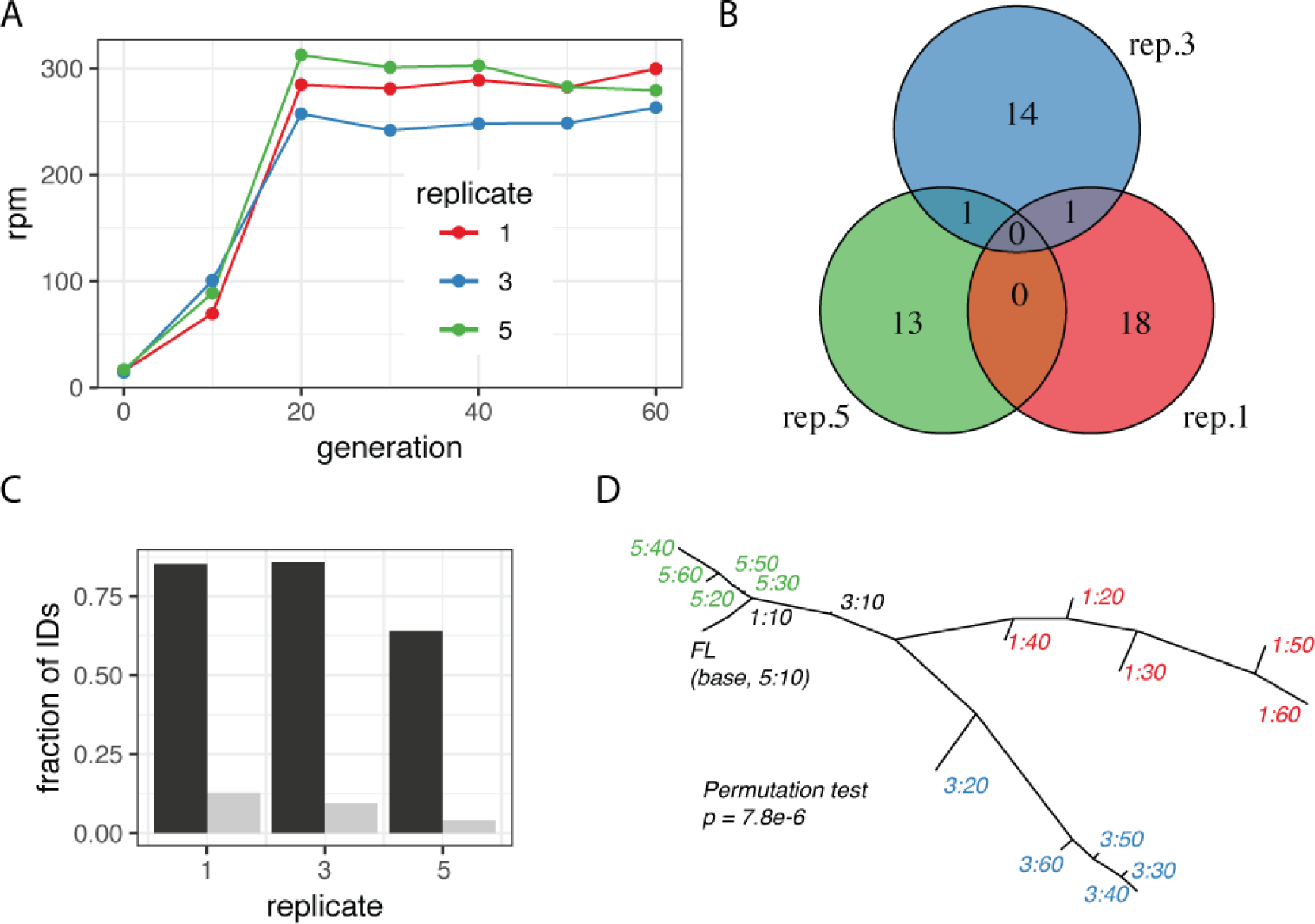
Each population invaded by the P-element receives a unique and persistent ID fingerprint. Data are from a replicated P-element invasion in experimental *D. simulans* populations [Kofler et al., 2018]. The populations were sequenced each 10^*th*^ generation as pools. A) In all three replicate populations, P-element copy numbers rapidly increased within the first 20 generations but no further increase was noted for the next 40 generations. B) Venn diagram of IDs found in different replicates (at any time point). Most IDs are replicate specific. C) IDs are more often recovered at different time points of the same replicate (black) than at any time point of a different replicate (grey). Hence, IDs persist within replicates. D) Unrooted tree constructed from the ID fingerprints of the populations. Once the invasions plateaued (≥20 generations), the populations (replicate:generation) lie on replicate specific clades. A population without IDs (FL) was included.

In summary, we showed that each independent P-element invasion generates an unique ID fingerprint that persists within populations. Furthermore, the similarity of ID fingerprints may be quantified with Jost’s D, which enables calculating a genetic distance that reflects the invasion history of samples. Samples from the same invasion have a small distance, whereas samples from independently invaded populations have larger distances.

Next, we asked whether IDs are useful markers for spatial population genetics. Using publicly available data, we were interest if IDs of the P-element allow to reproduce spatial signals found in previous works that relied on SNPs as markers. Some similarity in the spatial signal of SNPs and IDs may be expected, assuming that the migration pattern which shaped genome-wide polymorphism of SNPs also influenced the distribution of IDs. Bergland et al. [2014] found a latitudinal cline in sequenced *D. melanogaster* populations from the east coast of North America (“Bergland data”) and Kapun et al. [2018] found a longitudinal cline in populations from different locations in Europe (“DrosEU data”). In both data sets several IDs were found in more than one population (18 for Bergland; 124 for DrosEU; Fig. 3A; supplementary Fig. 1A). However, a large fraction of the IDs are solely found in a single population (321 in Bergland data; 572 in DrosEU data). Such a large fraction of population specific IDs may be a source of noise when assessing the genetic distance among populations based on IDs. In agreement with this, the spatial autocorrelation was significant for both data sets when population specific IDs where excluded from the analysis, but not when they were included (Mantel permutation test, 100, 000 permutations; geographic distance versus genetic distance based on ID fingerprints; Bergland data *p*_*incl*._ = 0.857, *p*_*excl*._ = 0.0103; DrosEU data *p*_*incl*._ = 0.0807, *p*_*excl*_ = 0.00001). Simulations confirmed that exclusion of these population specific IDs allows to estimate the genetic distance among populations more accurately (see below). Throughout this work, we thus ignored population specific IDs for estimating the relationship among populations with principal component analysis (PCA) and Jost’s D.

**Figure 3:**
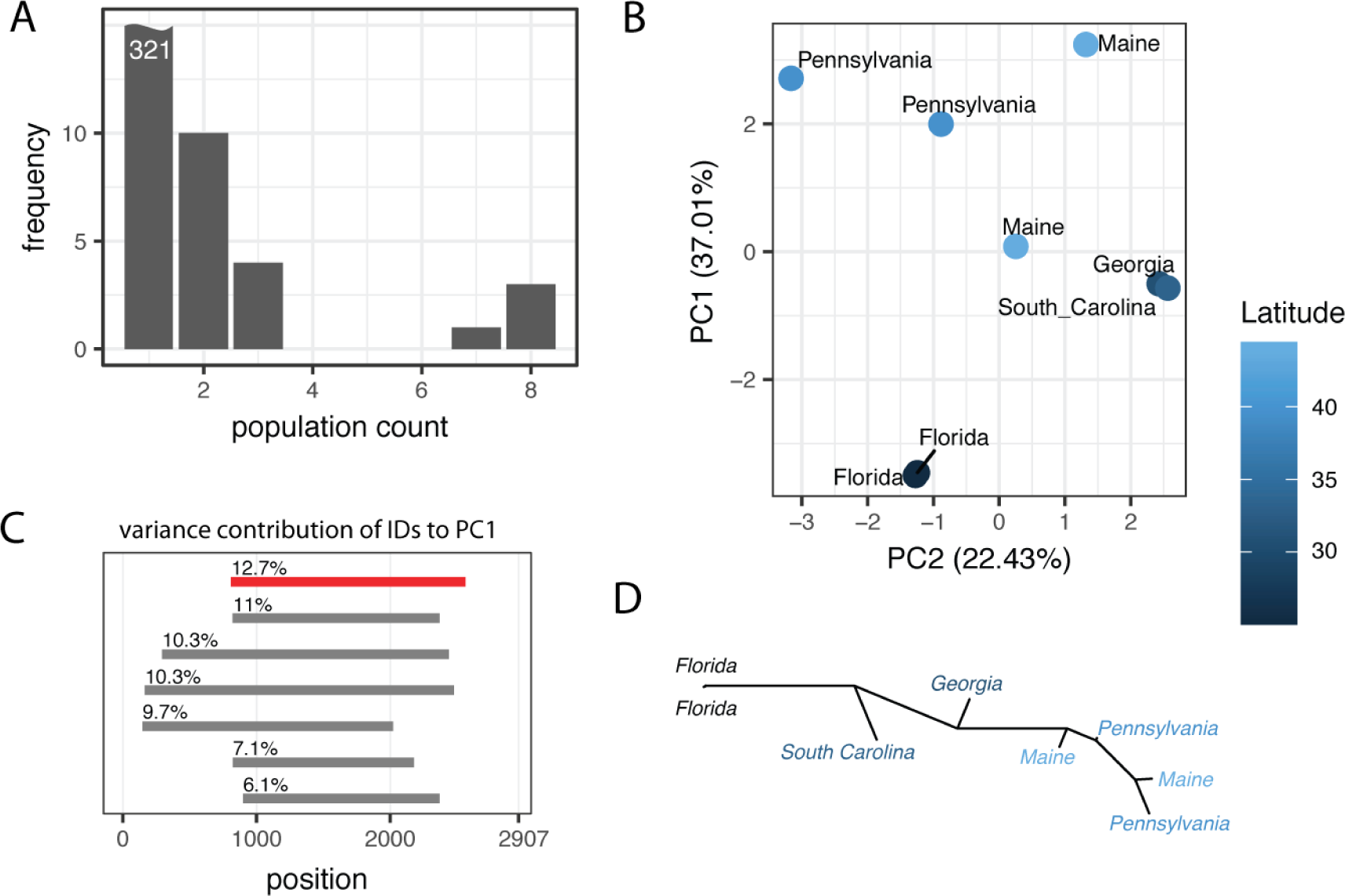
IDs of the P-element allow to reproduce the latitudinal cline found in *D. melanogaster* populations from the East Coast of the United States [Bergland et al., 2014]. A) Distribution of IDs in the populations. Most IDs (321) are only found in a single population while three are found in all 8 populations. B) PCA based on the frequency of IDs. PC1 is correlated with latitude. C) Variance contribution of the most important IDs to PC1. The start and the end position of IDs (line) as well as the variance contributions of IDs are shown. The KP-element is shown in red. D) Unrooted tree based on the genetic distance of the ID fingerprints. A latitudinal cline can be observed along the tree.

We performed a scaled PCA with the frequency of IDs found in the Bergland data (Fig. 3B). PC1 was significantly correlated with latitude (Spearman’s rank correlation *ρ* = 0.86, *p* = 0.0061). A weak correlation of PC1 with longitude was found (Spearman’s rank correlation *ρ* = 0.74, *p* = 0.036). No correlation with either longitude or latitude was found for PC2 (Spearman’s rank correlation; latitude *ρ* = −0.21, *p* = 0.62; longitude *ρ* = −0.01, *p* = 0.98). Since the KP-element, a distinct ID of the P-element where nucleotides 808-2560 are deleted, is widespread and highly abundant in worldwide populations of *D. melanogaster* [Black et al., 1987, Itoh and Boussy, 2002, Bergman et al., 2017], we asked whether the PCA results were mainly due to the KP-element. Investigating the contribution of the different IDs to PC1, we found that the KP-element indeed had the strongest influence to PC1, but several other IDs had very similar effects (Fig. 3C; KP-element shown in red). We also repeated the PCA without the KP-element and found that the significant correlation of PC1 with latitude could still be observed (Spearman’s rank correlation; *ρ* = 0.80, *p* = 0.017). The observed latitudinal cline is therefore due to frequency variation of multiple IDs. Finally, we used the genetic distance among ID fingerprints to generate an unrooted tree for the populations from the Bergland data (fig. 3D). This tree also reflects the latitudinal cline, where samples from the South (Florida) are located at one end and samples from the North (Maine) at the other end of the tree (Fig. 3D).

Next, we performed a scaled PCA with the frequency of IDs found in the DrosEU data (supplementary Fig. 1B). Both PC1 and PC2 showed only minor contributions to the total variance (*PC*1 = 9.51%, *PC*2 = 6.8%; supplementary Fig. 1B). However, both PC1 and PC2 were significantly correlated with longitude but not with latitude (Spearman’s rank correlation; longitude *ρ*_*PC1*_ = −0.81, *p*_*PC1*_ = 1.96*e -* 12, *ρ*_*PC2*_ = ™0.55, *p*_*PC2*_ = 5.41*e* − 05; latitude *ρ*_*PC1*_ = 0.21, *p*_*PC1*_ = 0.15, *ρ*_*PC2*_ = 0.003, *p*_*PC2*_ = 0.98;). Again, the KP-element only had a minor contribution to PC1 (supplementary Fig. 1C). A scaled PCA excluding the KP-element confirms the correlation of PC1 and PC2 with longitude but not with latitude (Spearman’s rank correlation; longitude *ρ*_*PC1*_ = −0.82, *p*_*PC1*_ = 1.20*e -* 12, *ρ*_*PC2*_ = −0.57, *p*_*PC2*_ = 2.14*e -* 05; latitude *ρ*_*PC1*_ = 0.22, *p*_*PC1*_ = 0.12, *ρ*_*PC2*_ = 0.05, *p*_*PC2*_ = 0.72;)

We conclude that IDs of the P-element allow to accurately reproduce spatial information described in previous works (longitudinal cline in DrosEU data and latitudinal cline in Bergland data). IDs are therefore useful markers for spatial population genetics.

To test whether IDs may be used to infer the invasion route of TEs we performed simulations. We aim to show i) that the fraction of ID insertions allows to estimate the direction of an invasion and ii) that successively invaded populations end up with similar ID fingerprints, which permits to infer the path of an invasion. The direction together with the path constitute the invasion route.

We generated a model that incorporates our current understanding of TE invasions and the biology of IDs. Initially we investigated the dynamics of TE invasions with IDs in a single population and later extend this model to multiple populations with migration. We simulated diploid organisms with 5 chromosomes of1 Mb, a recombination rate of 4 cM/Mb and piRNA clusters of 100 kb at one end of each chromosome (Fig 4A). We modeled the spread of a TE with a given transposition rate (*u*) in a population of size *N* = 1000 (Fig 4B). The TE invasions were launched by randomly distributing 300 FL elements in the genome (population frequency of insertions 1*/*2*N*). We assumed that IDs result from an interruption of sister chromatid mediated gap repair following excision of a FL element [Engels et al., 1990, Daniels and Chovnick, 1993, Gloor et al., 1991]. Hence, IDs are solely generated when the TE is active. A transposing FL element yields a novel ID with probability *c* (henceforth “conversion rate”; Fig 4B; top panel). For each novel ID we randomly picked two breakpoints (within positions 1 and 2907, i.e the length of the P-element). We assumed that FL and ID elments transpose at the same rate *u*, that every TE may solely harbour a single ID and that elements with an ID can not accumulate further IDs (Fig 4B). ID elements are assumed to be non-autonomous [Hua-Van et al., 2011]. Hence, mobilization of ID elements requires the presence of at least one FL insertion per diploid individual (Fig 4C). Finally, we simulated TE invasions under the trap model, which holds that the proliferation of an invading TE is stopped by TEs (FL or ID) that randomly jump into piRNA clusters [Kofler, 2019, Kelleher et al., 2018]. The TE is thus solely active in diploid individuals that do not carry a cluster insertion (Fig 4C, left). We assumed that both ID and FL insertions in piRNA clusters inactivate a TE, as previous works demonstrated that short stretches of homology between the cluster insertion and the TE sequence are sufficient for piRNA mediated silencing [Post et al., 2014].

**Figure 4:**
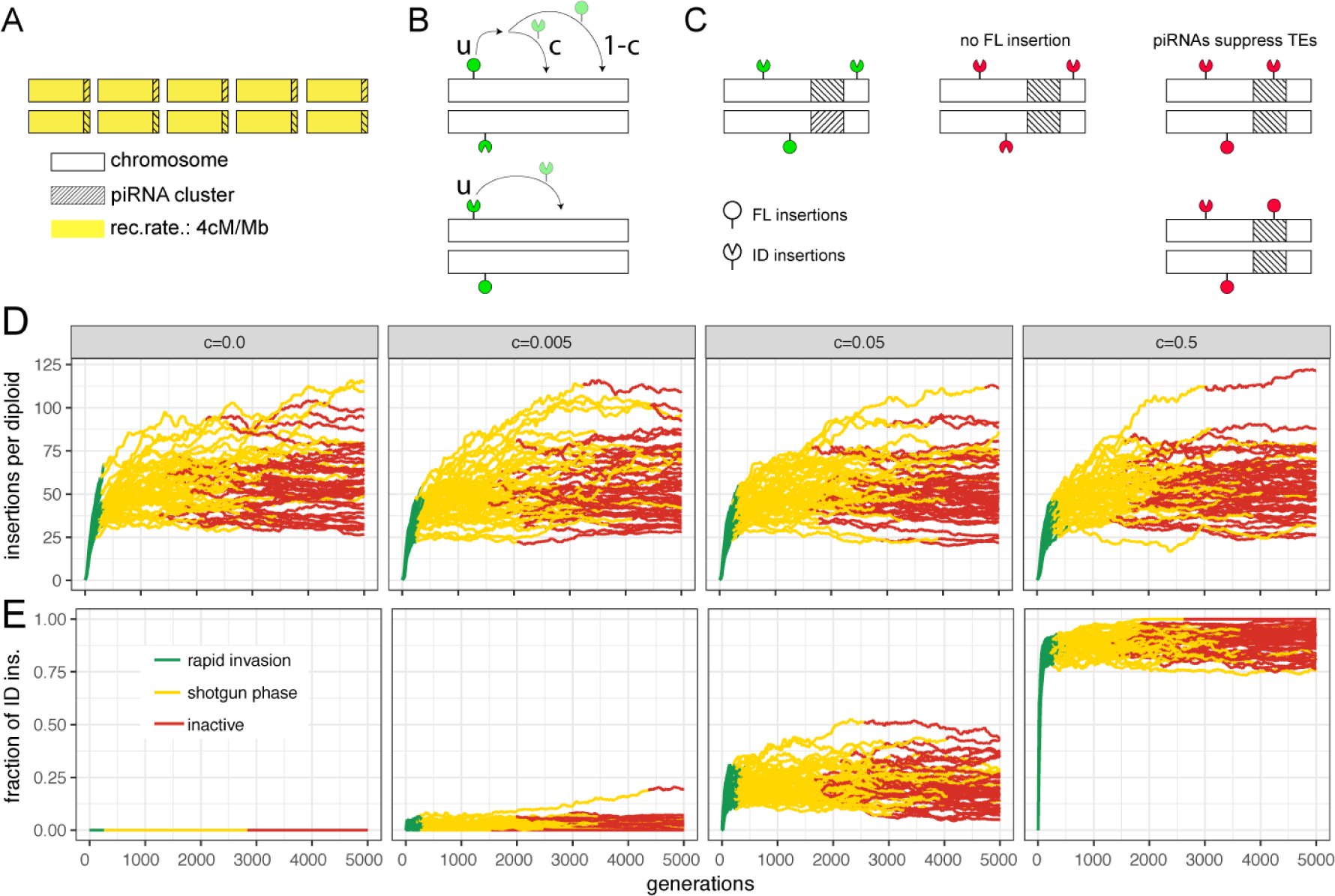
Dynamics of TE invasions with IDs. A) We simulated 5 chromosomes of 1 Mb and piRNA clusters of 100 kb at one end of each chromosome. Diploid individuals were simulated. B) While both FL and ID insertions are mobilized with a transposition rate *u*, solely FL elements may be converted into ID elements (with probability *c* per transposition event). C) In our model TEs are active (green) in individuals that have at least one FL insertion but no insertion (FL or ID) in a piRNA cluster. Absence of FL insertions or piRNA cluster insertions leads to inactive TEs (red). D) Influence of the conversion rate (*c*; top panel) on the TE abundance during an invasion. Data are shown for 50 replicates. The conversion rate has a negligible effect on the TE abundance. E) Fraction of ID insertions during a TE invasion. Note that the abundance of ID insertions solely increases at early stages of an invasion (green).

The dynamics of TE invasions with piRNA clusters have been explored before [Kelleher et al., 2018, Lu and Clark, 2009, Kofler, 2019]. In the absence of IDs (*c* = 0.0), we found the typical three phases of a TE invasion (Fig 4D; [Kofler, 2019]). First, the TE rapidly spreads in invaded populations (rapid invasion phase, green), then the TE is silenced by segregating cluster insertions (shotgun phase, yellow) and finally the TE is permanently inactivated by a fixed cluster insertion (inactive phase, red; Fig 4 [Kofler, 2019]). Interestingly, introducing IDs into our model (*c* > 0) had little influence on the invasion dynamics (Fig 4D). The length of the phases as well as the TE abundance were largely unaffected by IDs (supplementary Fig. 2). However, the conversion rate (*c*) had a profound effect on the abundance of ID elements (Fig 4E). The fraction of ID insertions increased rapidly during the early stages of an invasion (green) but plateaued when the TE was silenced (yellow, red; Fig 4E). The abundance of IDs at the plateau depended on the conversion rate (*c*), as for example 7% of TEs ended up with an ID with *c* = 0.005 and 25% with *c* = 0.05 (Fig 4E).

Importantly, the fraction of IDs did not increase further when the TE was silenced (Fig 4E; supplementary Fig. 3). This raises the question how worldwide populations may end up with vastly different numbers of IDs (assuming a constant *c* for a given TE family). For example, P-elements have many IDs in populations from North America but few in populations from Europe [Bergman et al., 2017].

Here we propose a simple explanation for the varying abundance of IDs observed in worldwide populations which, as a side effect, also highlights an approach for estimating the direction of an invasion. We suggest that migrants from an invaded population into a non-invaded population will introduce a sample of the insertions (FL and IDs) from the source population into the target population. This will trigger a novel invasion in the target population (Fig. 5A). Both FL and IDs introduced by the migrants will increase in copy numbers during the invasion. Additionally, however, novel IDs will be acquired due to the renewed TE activity (Fig. 5A). The target population will therefore, on the average, end up with more IDs than the source population (Fig. 5A). The fraction of IDs will increase in successively invaded populations and thus serves as a guide to the direction of an invasion. The origin of an invasion should be close to the population with the fewest IDs. We tested our hypothesis using a stepping stone model with 5 populations and a conversion rate of *c* = 0.025 (Fig. 5B). We initiated an invasion in the first population using 250 randomly distributed FL elements. We allowed TEs to invade a population for 300 generations before introducing the next migration event. This enabled populations to acquire distinct ID fingerprints (most TEs were silenced by generation 300). After 300 generations 100 migrants were moved from the invaded population into the next naive population, thus triggering a novel invasion (Fig. 5B). We repeated these migration events until all populations where infected by the TE (at generation 1200). Interestingly, the abundance of TEs remained constant in successively invaded populations (Fig. 5C; supplementary Fig. 4). However, the fraction of IDs significantly increased with each successively invaded population (Fig. 5D; supplementary Fig. 4). This confirms that the fraction of IDs serves as a rough guide to the direction of an invasion, where populations that were invaded first contain the fewest IDs.

**Figure 5:**
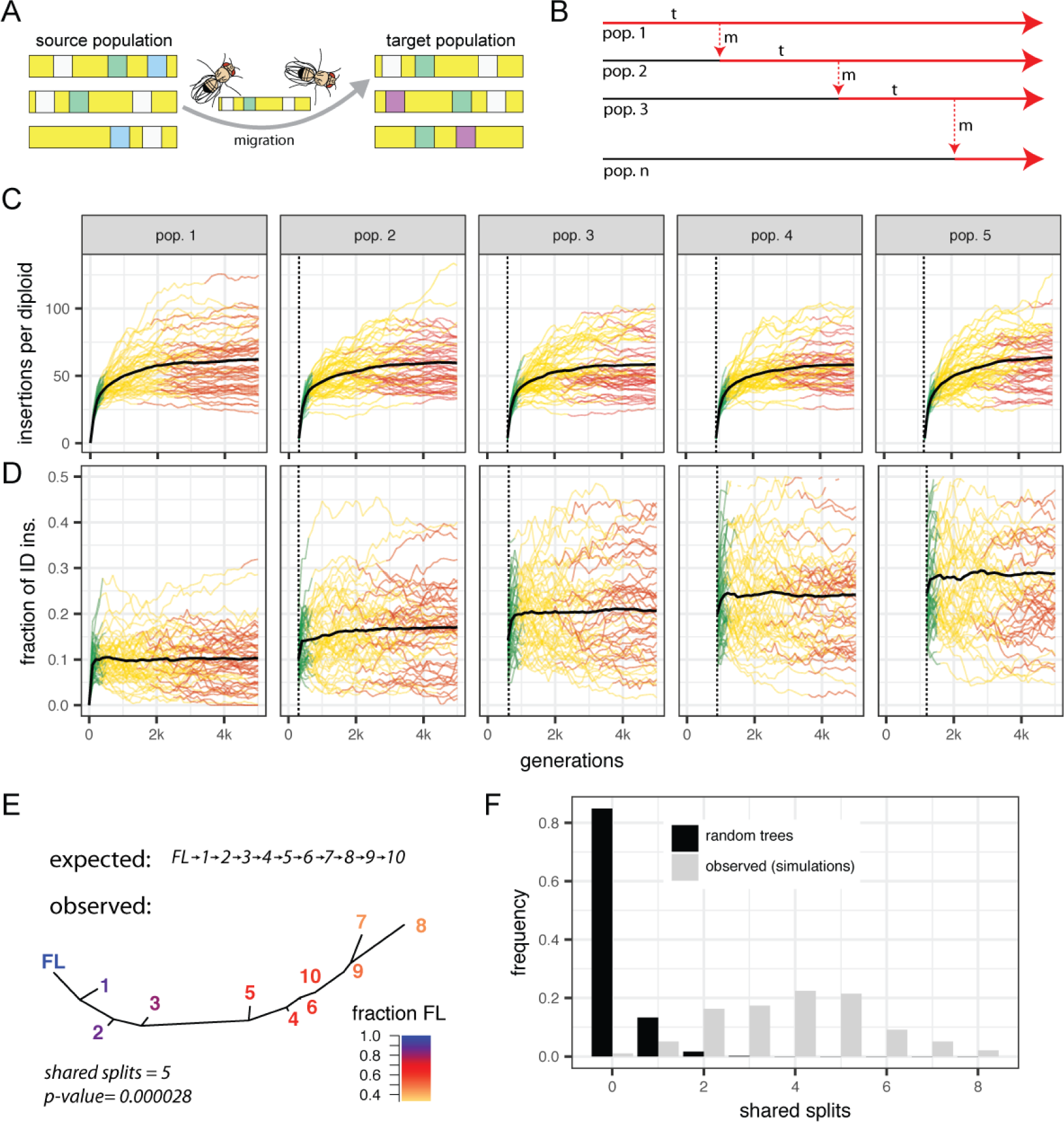
Model for reconstructing the spatial invasion history of TEs. A) Migrants from a source population carry some ID (colored boxes) and FL insertions (white boxes) to a target population, thereby triggering a novel invasion. During the invasion of the target populations, IDs introduced by migrants will be propagated (green) and novel IDs will be acquired (violet). B) Stepping stone model of successive TE invasions. After a TE invaded a population (*t* = 300), a number of migrants (red dashed line, *m* = 100) move to a naive population not having the TE (black line), triggering a new invasion (red line). C) TE abundance in successively invaded populations (top panel). The dashed lines indicate migration events that introduced the TE. The bold lines indicate the average over 50 simulations. D) Fraction of ID insertions in successively invaded populations. Note that the fraction of IDs increases with each invaded population. E) Example of a reconstructed TE invasion. Based on ID fingerprints, a distance matrix and a tree was constructed. Numbers refer to the order of the invaded populations. As origin of the invasion a FL insertion was included (FL). Note that the fraction of FL insertions decreases with each invaded population. The tree shares 5 splits with the expected one, which is highly significant. F) Accuracy of reconstructed TE invasions for 100,000 random trees and 100 trees derived from our simulations. In this scenario the highest possible number of shared splits is 8.

Next, we tested if we may infer the path of an invasion based on the similarity of ID fingerprints among populations. Using the stepping stone model introduced above, we simulated TE invasions in 10 successive populations. We used Jost’s D to estimate the similarity of ID fingerprints among populations at generation 3000, i.e. after all 10 populations were invaded (the last migration event occurred at generation 2700). This results in a matrix of pairwise distances among populations. To mark the origin of the invasion, we included an artificial population consisting solely of FL elements into this distance matrix. Finally, we inferred trees from the distance matrices and compared them to the expected tree (expected invasion route: FL →1 → … → 10). We assessed the accuracy of the inferred trees using the number of splits shared between the observed and the expected tree as quality metric (supplementary Fig. 5). A split, or bifurcation, is the smallest information unit of unrooted trees and the number of splits shared between two trees may be used as a similarity metric when solely the topology of trees is considered (i.e. the branch length is ignored). In our scenario the largest possible number of shared splits is 8. As example, a tree with 5 shared splits already allows to fairly accurately infer the invasion route (Fig. 5E). To derive the null expectation, we computed the shared splits for 100,000 random trees having the same tips as the expected trees (Fig. 5F). On average, random trees have 0.17 shared splits with the expected tree. Our approach allows reconstructing invasion trees with an average precision of 3.9 shared splits (Fig. 5E; 100 simulations), which is significantly better than random expectations (Wilcoxon rank sum test; *W* = 96819120, *p* < 2.2*e* − 16). Since genetic drift may distort ID fingerprints over time, we asked how long invasion routes could be estimated. We thus inferred invasion routes at different time points after invasion of all 10 populations (≥3000 generations). Invasion routes may be traced for hundreds, possibly even thousands, of generations after the spread of the TE, with only a small loss of accuracy with time (supplementary Fig. 6).

In summary, we showed that it is feasible to infer the invasion route of DNA transposons using IDs as markers. The similarity of ID fingerprint provides cues about the invasion path and the abundance of FL insertions acts as a rough guide to the direction of an invasion.

Next, we applied our approach to worldwide populations of *D. melanogaster* to test if the reconstructed invasion route of the P-element agrees with previous work. The global P-element invasion in *D. melanogaster* likely started in South America following a horizontal transfer from *D. willistoni* [Daniels and Chovnick, 1993]. The P-element first spread within South and North American populations, and later in European and African populations [Anxolabéhère et al., 1988]. It is, however, not clear, if European or African populations were invaded first. In Africa the P-element was first observed in the South, whereas in Europe the P-element was first found in France [Anxolabéhère et al., 1985, Anxolabéhère et al., 1988]. Starting from France, the P-element spread to Spain and towards the East of Europe [Anxolabéhère et al., 1985, Anxolabéhère et al., 1988].

To infer the invasion route of the P-element, we relied on the Bergland and DrosEU data mentioned above [Kapun et al., 2018, Bergland et al., 2014], as well as the Dros-RTEC data from North America [Machado et al., 2018] and the DPGP2/3 data from Africa [Lack et al., 2015]. To avoid biases we discarded reads smaller than 90bp and trimmed all reads to a size of 100bp. For an overview of the abundance of FL and ID P-elements in these populations, see supplementary Fig. 7.

We estimated the similarity of ID fingerprints among populations using Jost’s D and visualized the resulting distance matrix with a multidimensional scaling (MDS) plot and a tree (Fig. 6). Neighboring samples in the MDS plot have similar ID fingerprints. An artificial population containing solely FL elements was added to mark the origin of the invasion (Fig. 6).

**Figure 6:**
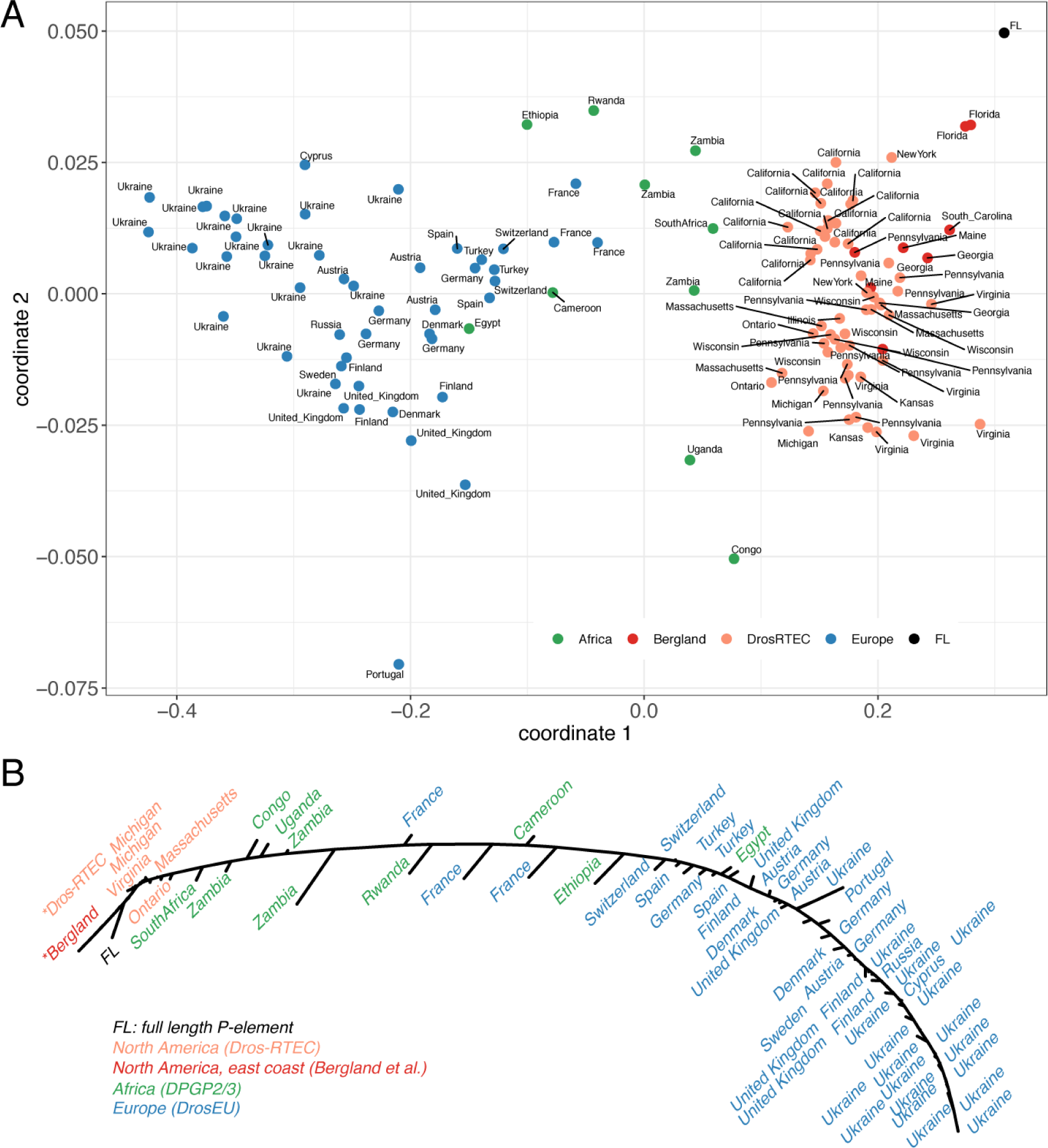
Reconstructed invasion route of the P-element in worldwide populations of *D. melanogaster*. A) Multidimensional scaling (MDS) plot based on the similarity of ID fingerprints among populations. Note that African samples clusters between North American and European samples. B) Tree showing the invasion route of the P-element. Starting from the FL insertion, the P-element first invaded North America, then spread from southern to northern Africa and finally invaded Europe from West to East.

Based on our approach we suggest that the P-element invaded populations from Florida and California first and then spread into other North American populations (Fig. 6. Interestingly, our data suggest that the P-element invaded African populations prior to European ones (Fig. 6; supplementary Fig. 8). Within Africa the P-element spread from South to North, until it eventually invaded Europe (Fig. 6). Starting from France, the P-element spread to Spain and towards the East of Europe. Lastly, populations from Ukraine were invaded (Fig. 6).

The invasion route of the P-element inferred by our approach is remarkably similar to the route proposed by previous works, which relied on fly strains sampled over the course of decades from different geographic locations [Anxolabéhère et al., 1985, Anxolabéhère et al., 1988].

Finally, we applied our method to a TE with a controversial invasion history, i.e. hobo in *D. melanogaster* (Fig. 7). Similarly to the P-element, hobo invaded worldwide *D. melanogaster* populations within the last 100 years [Daniels et al., 1990, Pascua and Periquet, 1991]. However, the hobo invasion likely predates the P-element invasion [Periquet et al., 1989, Pascua and Periquet, 1991].

**Figure 7:**
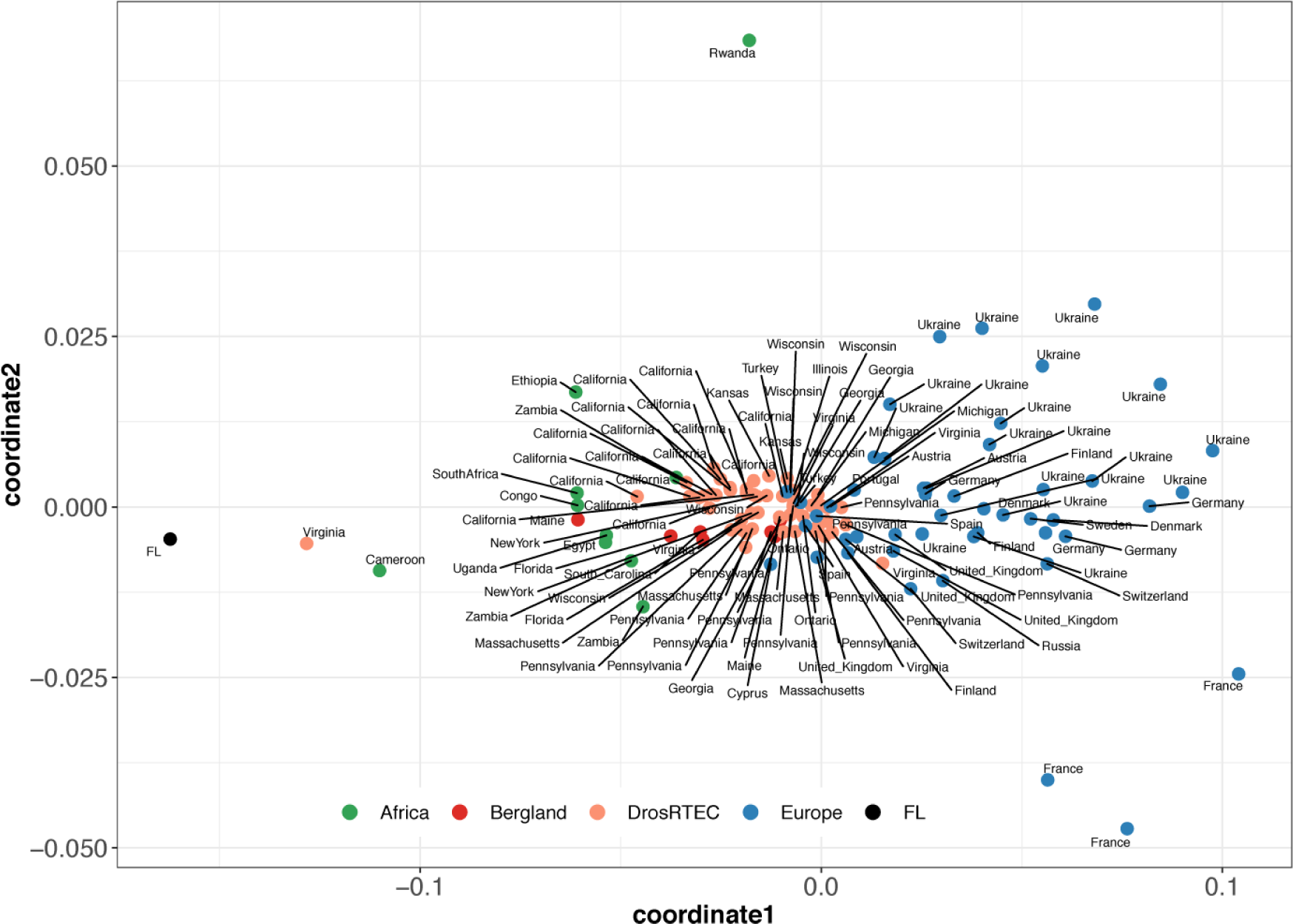
Invasion route of the hobo element in worldwide populations of *D. melanogaster*. The MDS plot is based on the similarity of ID fingerprints among populations. We suggest that hobo invaded African populations first, then spread to North America and finally invaded European populations.

Many strains collected in North America before 1955 contained hobo sequences, whereas few of the European strains collected before 1955 contained hobo sequences [Periquet et al., 1989]. It was thus suggested that the hobo invasion started in North America [Periquet et al., 1989]. Alternatively, it was suggested that Europe could be the origin of the hobo invasion since the diversity of internal tandem repeats is highest in European populations [Bonnivard et al., 2000]. Finally, populations from Kenya showed the highest hobo activity, raising the possibility that hobo spread from Africa to other continents [Bonnivard et al., 2000].

Based on our approach, we suggest that hobo invaded populations from Africa first, then spread to North America and finally invaded European populations (Fig. 7). Similarly to the P-element, populations from Ukraine were invaded last (Fig. 7). The route inferred by our approach is thus in agreement with the proposed invasion of North American populations prior to European ones [Periquet et al., 1989] and an origin of the hobo invasion in Africa [Bonnivard et al., 2000], but not with an origin of the invasion in Europe [Bonnivard et al., 2000].

The reconstruction of the P-element and hobo invasion routes with our approach confirms that it is feasible to infer the invasion route of DNA transposons using extant population samples.

## Discussion

Here we propose that IDs allow to trace the invasion route of DNA transposons. This is made possible by an interesting property of IDs of DNA transposons: they emerge at a high rate solely when the TE is active [Engels et al., 1990, Daniels and Chovnick, 1993, Gloor et al., 1991]. As a consequence, each independently invaded population receives a unique ID fingerprint, which remains recognizable over multiple generations (Fig. 2). Furthermore, we found that IDs of DNA transposons are useful markers for spatial population genetics. For example, IDs of the P-element support the previously described clines in *D. melanogaster* populations based on genome wide SNP data [Machado et al., 2018, Kapun et al., 2018]. It was, however, not necessarily expected that IDs and SNPs show a similar spatial signal. A TE invasion could have also taken a route that differs from the migration paths that shaped the polymorphism pattern of SNPs. It is, however, plausible that dominant migration patterns also influence invasion routes. Furthermore, migration following the invasion may also have influenced the distribution of IDs (see below). Interestingly, clines in populations from Europe and North America could not be observed when other properties of TEs, such as TE abundance and TE diversity were investigated [Adrion et al., 2019, Lerat et al., 2019]. We thus argue that IDs are sensitive markers that allow to pick up spatial signals that are otherwise not found with TEs.

Surprisingly, many IDs were solely found in a single population (Fig. 3; supplementary Fig. 1). In Europe, for example, 527 out of 696 IDs were specific to a single population (supplementary Fig. 1). The reason for this larger number of population specific IDs may partly be technical. As the majority of IDs occur at a very low frequency in populations, only a small number of IDs will be present in a sample and in the sequencing data. This could lead to a low overlap of IDs among populations, even when these populations have very similar ID compositions. Similarly, migrants between populations will only carry a small sample of the IDs present in populations. Homogenizing the distribution of IDs among populations may thus be a very slow process. Alternatively, the P-element could still be active in some populations. Novel populations specific IDs may yet emerge in these populations. However, we deem this scenario unlikely. In a previous work we found that a piRNA based defence mechanism against an invading P-element rapidly emerged in *D. simulans* [Kofler et al., 2018]. Within a mere 20 generations P-element proliferation was stopped in all investigated replicates [Kofler et al., 2018]. The P-element invaded worldwide *D. melanogaster* populations several hundred generations ago [1950-1980 with 15 generations per year [Anxolabéhère et al., 1988, Pool, 2015]]. This should be sufficient time for establishing a piRNA based defence against the P-element. Solely an equilibrium state, termed transposition-selection-cluster balance (TSC balance), may account for persistent TE activity in the face of a piRNA based defence mechanism [Kofler, 2019]. In this scenario negative selection against all TE insertions, including piRNA cluster insertions, prevents permanent inactivation of a TE [Kofler, 2019]. However, TSC balance was so far solely demonstrated in a theoretical model and it is unclear if this equilibrium state can be found in natural populations.

Our approach for reconstructing the invasion route of DNA transposons rests on two assumptions. First, that the abundance of IDs in populations provides cues about the direction of an invasion, as the fraction of IDs increases in successively invaded populations. Second, that we may infer the path of an invasion by the similarity of ID fingerprints among populations. Our hypothesis that early invaded populations end up with the most FL elements, may be regarded as somewhat counterintuitive. It could be argued that early invaded populations should have the fewest FL elements, since the TE was active the longest in these populations which should have allowed them to accumulated most IDs [e.g. Bergman et al., 2017]. However, this view does not consider that TE invasions are rapidly silenced by segregating piRNA cluster insertions [Kofler, 2019, Kofler et al., 2018, Kelleher et al., 2018] and that no further increase in the fraction of IDs is expected once the TE is silenced (supplementary Fig. 3). The fraction of IDs may solely increase if a silenced TE is reactivated by some means. Here we propose that a TE may be reactivated by migrants that trigger a new invasion in a naive population (Fig. 5). During this invasion, IDs introduced by migrants will be amplified and novel IDs will emerge. As a consequence, the fraction of IDs will increase in successively invaded populations (supplementary Fig. 4). Our model accounts for the striking differences in the abundance of P-element IDs between populations from Europe and North America [Black et al., 1987, Bergman et al., 2017]. Since European populations have more IDs than North American ones, our model predicts that Europe was invaded after North America, which is in agreement with previous works [Anxolabéhère et al., 1988]. Alternatively, it was proposed that North American populations may be able to control IDs more efficiently than European populations [Bergman et al., 2017]. This model has two problems: it requires a hitherto unknown mechanism for controlling IDs; and it does not explain how differences in the regulation of IDs among continents may emerge. Our model provides a simpler explanation.

We propose that similarities of ID fingerprints among populations allow to trace the path of an invasion. The accuracy of the inferred path will depend on many factors such as the used distance metric, the migration rate, the migration pattern, and the conversion rate. Based on our simulations we evaluated the performance of some distance metrics for estimating the similarity of ID fingerprints among populations. The difference in the fraction of FL insertions had the worst performance (shared splits = 2.4), followed by Jost’s D (shared splits = 3.4; supplementary Fig. 9)). Removing IDs solely found in a single population significantly increased the accuracy of Jost’s D (mean shared splits = 4.0; Wilcoxon rank sum test *p* = 0.013; supplementary Fig. 9). Finally, the inverse of the number of IDs shared between two populations (1*/shared*) performed best (shared splits = 5.3; supplementary Fig. 9). This approach may, however, not be suitable for real data as the number of shared IDs will be highly susceptible to differences in sampling and sequencing depths among samples (in our simulations all IDs are known).

The conversion rate *c* is a crucial parameter. If the conversion rate is low, few IDs will emerge and tracing the invasion will not be feasible. Our simulations suggest that conversion rates *c* > 0.01 are necessary to trace invasions with reasonable accuracy (supplementary Fig. 10). During a P-element invasion in *D. simulans*, the fraction of IDs in populations plateaued at about 13% [Kofler et al., 2018]. We estimate that this corresponds to a conversion rate of *c* ≈ 0.03 (interpolating the plateauing level from our simulations, Fig. 5). Thus, the conversion rate of the P-element, at least in *D. simulans*, is well within the parameter range resulting in accurate invasion paths (supplementary Fig. 10). It is, however, not clear if conversion rates of other DNA transposons are also sufficiently high. Nevertheless, we found that other DNA transposons also carry large amounts of IDs (supplementary Fig. 11). Such families are attractive future targets for tracing invasions with our approach. Surprisingly, many LTR and non-LTR transposons also carry substantial amounts of IDs (supplementary Fig. 12). It remains to be seen whether these IDs could be used for reconstructing TE invasions.

In our model we assumed that both FL and ID elements are mobilized at the same rate (i.e. in the presence of an autonomous FL element). It was, however, suggested that IDs may be mobilized at a higher rate than FL elements [Itoh et al., 2007, Kofler et al., 2018]. Preferential mobilization of IDs greatly enhances the accuracy of our approach (supplementary Fig. 13). For example, if IDs are 25% more readily mobilized than FL elements, the accuracy of the reconstructed invasion route increased by 50% (supplementary Fig. 13).

The migration rate also influences the accuracy of our approach. If the number of migrants is small, few IDs will be carried to the target population, and thus the similarity of ID fingerprints among populations will be too low to for reconstructing invasion routes. In our simulations about 50 migrants were necessary to trace invasions with reasonable accuracy (supplementary Fig. 14). Interestingly, larger numbers of migrants (*>* 50) had only a minor effect on the accuracy of our approach (supplementary Fig. 14). Migration pattern also influence the accuracy of our approach. Initially we simulated unidirectional and unique migration events from an invaded source population into a naive target population, thus triggering a novel invasion in the target population (Fig. 5B). With this approach we ensured that the inferred invasion route is solely based on the migration events that triggered the invasion and not on migration events following the invasion. However, in natural populations migration between neighboring populations will likely occur more regularly and in both directions. Interestingly, bidirectional and recurrent migration (100 migrants in both directions every 300^*th*^ generation) increased the accuracy of our approach by 41% (from 3.9 to 5.5 shared splits at generation 3000; supplementary Fig. 15). Furthermore, invasions could be traced with a high precision for hundreds of generations after the spread of the TE (*>* 3000 generation; supplementary Fig. 15). Recurrent migration likely increases the similarity of ID fingerprints between neighboring populations, which facilitates reconstructing the invasion route. This conclusion, however, rests on a crucial assumption, i.e. that migration routes are fairly stable over time. If migration routes change after the spread of a TE, recurrent migration will decrease the accuracy of our approach. If the novel migration routes persist, the inferred invasion routes will eventually reflect the novel migration routes rather than the invasion routes. Our method may also be inaccurate if invasion routes deviate from the major migration routes. It is however unclear if this scenario is important, as major migration routes will usually carry more TE bearing migrants to naive host populations than minor ones. So far we introduced migration events at every 300^*th*^ generation. With this approach we allowed TEs to completely invade a population before introducing the next migration event from the most recently invaded population to the next naive population. Within these 300 generations, TEs will mostly be silenced by segregating piRNA cluster insertions and populations will have acquired fairly stable ID fingerprints (Fig. 2D). It may, however, be argued that such a long pause between migration events is an implausible model for natural populations. With natural TE invasions the time required to silence a TE could actually be substantially shorter. For example, the P-element invasion in experimental *D. simulans* populations was silenced within a mere 20 generations [Kofler et al., 2018]. The reason for this discrepancy between real and simulated data, is however not clear (the transposition rate was similar: *u* = 0.1). We speculate that an insertion bias into the piRNA cluster, as described for the P-element, could accelerate silencing of the TE [Kofler et al., 2018, Karpen and Spradling, 1992, Zhang and Kelleher, 2019]. Paramutations, which convert euchromatic TE insertions into piRNA producing loci could also speed up silencing of an invasion [de Vanssay et al., 2012, Mohn et al., 2014]. Hence, natural populations may acquire stable ID fingerprints much faster than our simulated populations. Nevertheless, we found that our approach also works when populations do not have sufficient time to establish stable ID fingerprints. With a single bidirectional and recurrent migrant between neighboring populations at each generation, the invasion route can still be inferred, albeit with a reduced accuracy (supplementary Fig. 16). It is conceivable that other factors, not evaluated here, also influence the accuracy of our approach, such as the population size, the number of sampled individuals, and the size of piRNA clusters.

To trace the invasion of the P-element and hobo we applied our approach to available sequencing data from *D. melanogaster* populations. The inferred invasion routes fit remarkably well with previous works [Anxolabéhère et al., 1985, Anxolabéhère et al., 1988, Periquet et al., 1989, Bonnivard et al., 2000]. Sufficient data were, however, solely available for populations from North America, Europe and Africa. For the P-element, it would be especially interesting to extend our approach to populations from Central Asia and South America. Since South America is the likely origin 0of the P-element invasion [Daniels et al., 1990], populations from South American ought to have - according to our model - the most FL elements. Testing this hypothesis would be important.

Central Asian populations are interesting because they may not yet have the P-element. We found that the fraction of FL elements decreased in Europe from West to East, where very few FL elements were found in Ukrainian populations (supplementary Fig. 7). However, P-element activity requires autonomous FL insertions. The continuous dilution of FL elements from West to East raises the intriguing possibility that the P-element invasion may have run out of autonomous FL insertions in the East. Invasions of some DNA transposons may thus have a maximum range, beyond which the TE can not spread due to lack of FL elements. It would thus be interesting to test if populations East of Ukraine contain P-elements. Concerning hobo it would also be important to analyze populations from different geographic regions. In the MDS plot the FL element marking the origin of the invasion is quite distant from the population samples (Fig. 7), which indicates that the hobo invasion may have started in a geographic region that was not analyzed in this work.

We showed that an approach based on sequencing extant populations as pools, taking ID fingerprints of TEs with DeviaTE and computing the similarity of ID fingerprints with Jost’s D allows to reconstruct TE invasions with reasonable accuracy. It is possible that future advances in sequencing technology (read length, accuracy, throughput) and algorithms for identifying IDs will enhance the accuracy of our approach. As a major advantage, our approach allows to trace TE invasions using samples from extant populations. Previous works that aimed to reconstruct invasion routes of TEs relied on strains collected from different geographic origins over decades [Anxolabéhère et al., 1985, Anxolabéhère et al., 1988, Periquet et al., 1989]. Such time series data from different regions are, however, solely available for few species. Alternatively, sampling of time series data could be specifically initiated to trace an ongoing TE invasion. However, TE invasions are rare and most TE invasions are discovered only when most populations are already infected [e.g. Kofler et al., 2015, Pascua and Periquet, 1991, Jordan et al., 1999, Bucheton et al., 1992]. Once a TE invasion is detected, it is thus usually already too late to start sampling time series data. Due to these enormous difficulties, we only have very sparse knowledge about invasion routes of TEs. Our approach may change this dismal situation. As a disadvantage, however, our approach does not allow to estimate the time span in which the invasion happened, information that is available with time series data. Our approach requires sequences of TEs of interest and sequencing data of populations (or strains) from different geographic locations. Therefore, our approach could be used with model as well as non-model organisms. Due to the efforts of international consortia and individual research groups, population data from different geographic regions will increasingly become available for many diverse species [Kapun et al., 2018, Machado et al., 2018, Alonso-Blanco et al., 2016, Telenti et al., 2016]. Hence, our approach may increasingly be extended to different TE families in many diverse species. An enhanced understanding of the course of TE invasions will shed light on the biology and life cycle of TEs, the origin of TE invasions, migration patterns, and possibly even the ecology of species.

## Material and Methods

### Publicly available data

To analyse IDs during a P-element invasion in *D. simulans* we used the data of Kofler et al. [2018]. Further, we analysed Pool-Seq data of natural *D. melanogaster* populations sampled from Europe (DrosEU [Kapun et al., 2018]), North America (Dros-RTEC [Machado et al., 2018]), the East Coast of North America [Bergland et al., 2014] and Africa (DPGP2/3 [Lack et al., 2015]). For the data of Bergland et al. [2014] we used the samples with the most consistent sampling time (Linvilla, 2009: SRR1525768,SRR1525769). For DPGP2/3 data we only used samples of complete genomes having a minimum read length of 100bp and more than 1,000,000 reads. Solely sequences of individual strains were available for the DPGP2/3 data. Therefore, we artificially pooled them by sampling equal numbers of reads from strains with identical geographic origins. For an overview of all samples used in this work see supplementary Table 1.

### Analysis of data

All genomic data were downloaded using SRA-Tools (http://ncbi.github.io/sra-tools/) and aligned to a reference consisting of the consensus sequences of TEs from *D. melanogaster* (v9.44; https://github.com/bergmanlab/transposons; [Quesneville et al., 2005]) and sequences of the single copy genes *Rhino, RpL32* and *traffic jam*. To avoid heterogeneous read lengths, we trimmed all reads to a length of 100bp. Reads were mapped with bwasw [Li and Durbin, 2009] and sorted with samtools [Li et al., 2009]. Subsequently, the position and the frequency of IDs was estimated with DeviaTE [Weilguny and Kofler, 2019]. A custom script (*dm-deviate*.*py*) was used to estimate the genetic distance among samples. We treated each ID and the FL insertion as an allele of a TE and used Jost’s D [Jost, 2008] to compute the genetic distance among samples. The frequency of the FL insertion was computed as 1 −∑*f*_*i*_, where *f*_*i*_ are the frequencies of all IDs. Since alignments of reads with indels are difficult, we allowed for a tolerance of 3bp in the position of breakpoints. We required a minimum support of 2 reads for each ID in the data of [Kofler et al., 2018] and 3 reads in data from natural *D. melanogaster* populations. Finally, IDs solely occurring in a single population were ignored for PCA and estimates of Jost’s D. Distance matrices based on Jost’s D were used to generate MDS plots and unrooted trees with the bionj algorithm [Gascuel, 1997] implemented in the R package ape [Paradis et al., 2004]. PCA was performed with the frequency of IDs using R (prcomp; [R Core Team, 2014]). The geographic distance among samples was estimated with the R package geosphere [Hijmans, 2017] and the Mantel test for estimating the correlation of distance matrices was performed with the R package cultevo [Stadler, 2018].

### Simulations

Simulations were performed with a modified version of the Java tool Invade [Kofler, 2019]. The adapted tool invade-td (*t* runcations *d* eme) performs individual based forward simulations of TE invasions with piRNA clusters and IDs under a one dimensional stepping-stone model [Kimura and Weiss, 1964]. At each generation the tool performs the following steps in the given order 1) mate pairs are formed based on the fitness of the individuals, 2) haploid gametes are generated based on the recombination map, 3) novel FL and ID insertions are introduced, 4) zygotes are formed, 5) piRNA cluster insertions are counted, 6) the fitness of the individuals is computed, 7) migrants are exchanged between adjacent populations (optional) and 8) the output is produced (optional). Novel IDs emerged during the transposition phase (3) with probability *c* per transposing FL element. Two random breakpoints were chosen for each novel ID (between positions 1 and 2907, i.e. the length of the P-element). If not mentioned otherwise, we simulated a genome with five chromosomes of 1Mb (–genome Mb:1,1,1,1,1), a recombination rate of 4 cM/Mb (–rr cM Mb:4,4,4,4,4), piRNA clusters at the end of each chromosome (–cluster kb:100,100,100,100,100), a transposition rate of 0.1 (–u 0.1), neutral TE insertions (–x 0.0), a population size of 1000 (–N 1000) and equal mobilization of FL and ID insertions (–fl id 0.5). TE invasions were launched by introducing 250 to 300 randomly distributed FL insertions into the first population (frequency of *f* = 1*/*2 * *N*). The position and the frequency of the IDs (i.e. the ID fingerprint) was recorded in the output. We used a custom script to compute the genetic distance among simulated populations based on the ID fingerprints (*dm-simulations*.*py*). To mirror the treatment of samples from natural populations, we ignored IDs solely occurring in a single population. Trees were again generated with the bionj algorithm and the number of splits shared between the expected and observed trees were computed with the R package ape (comparePhylo; [Paradis et al., 2004]). To obtain the expected number of shared splits under the null model, we simulated 1 million random trees with the same tips as the expected tree (rtree; [Paradis et al., 2004]).

## Data availability

The simulation tool invade-td.jar (https://sourceforge.net/projects/invade/) as well as the scripts used in this work (https://sourceforge.net/p/te-tools; folder reconstruct) are freely available.

## Author contributions

RK and LW conceived the work. CV, LW, RK and DS analyzed the data. RK performed the simulations. RK wrote the manuscript with input from all authors.

## Acknowledgements

We thank Carolin Kosiol and Rui Borges for helpful comments about phylogenetic trees and Claus Vogel for feedback. We thank all members of the Institute of Population Genetics for feedback and support. This work was supported by an Austrian Science Foundation (FWF) grant (P30036-B25) to RK.

## References

J. R. Adrion, D. J. Begun, and M. W. Hahn. Patterns of transposable element variation and clinality in Drosophila. Molecular Ecology, 28(6):1523–1536, 2019.

Alonso-Blanco, J. Andrade, C. Becker, F. Bemm, J. Bergelson, K. M. Borgwardt, J. Cao, E. Chae, T. M. Dezwaan, W. Ding, et al. 1,135 genomes reveal the global pattern of polymorphism in *Arabidopsis thaliana*. Cell, 166(2):481–491, 2016.

Anxolabéhère, D. Nouaud, G. Périquet, and P. Tchen. P-element distribution in Eurasian populations of *Drosophila melanogaster*: a genetic and molecular analysis. Proceedings of the National Academy of Sciences, 82(16):5418–5422, 1985.

D. Anxolabéhère, M. G. Kidwell, and G. Periquet. Molecular characteristics of diverse populations are consistent with the hypothesis of a recent invasion of *Drosophila melanogaster* by mobile P elements. Molecular biology and evolution, 5(3):252–69, 1988.

A. O. Bergland, E. L. Behrman, K. R. O’Brien, P. S. Schmidt, and D. A. Petrov. Genomic Evidence of Rapid and Stable Adaptive Oscillations over Seasonal Time Scales in Drosophila. PLoS Genetics, 10(11): e1004775, 2014.

C. M. Bergman, H. Quesneville, D. Anxolabéhère, and M. Ashburner. Recurrent insertion and duplication generate networks of transposable element sequences in the Drosophila melanogaster genome. Genome biology, 7(11):R112, 2006.

C. M. Bergman, S. Han, M. G. Nelson, V. Bondarenko, and I. Kozeretska. Genomic analysis of P elements in natural populations of *Drosophila melanogaster*. PeerJ, 5:e3824, 2017.

Biémont and C. Vieira. Genetics: junk DNA as an evolutionary force. Nature, 443(7111):521–524, 2006.

M. Black, M. S. Jackson, M. G. Kidwell, and G. A. Dover. KP elements repress P-induced hybrid dysgenesis in *Drosophila melanogaster*. The EMBO journal, 6(13):4125–35, 1987.

J. P. Blumenstiel, X. Chen, M. He, and C. M. Bergman. An Age-of-Allele Test of Neutrality for Transposable Element Insertions. Genetics, 196(2):523–538, 2014.

E. Bonnivard, C. Bazin, B. Denis, and D. Higuet. A scenario for the hobo transposable element invasion, deduced from the structure of natural populations of *Drosophila melanogaster* using tandem TPE repeats. Genetical Research, 75(1):13–23, 2000.

J. Brennecke, A. A. Aravin, A. Stark, M. Dus, M. Kellis, R. Sachidanandam, and G. J. Hannon. Discrete small RNA-generating loci as master regulators of transposon activity in Drosophila. Cell, 128(6):1089–1103, 2007.

A. Bucheton, C. Vaury, M. C. Chaboissier, P. Abad, A. Pélisson, and M. Simonelig. I elements and the *Drosophila* genome. Genetica, 86(1-3):175–90, 1992.

A. Burt and R. Trivers. Genes in conflict: the biology of selfish genetic elements. Belknap Press, 2008.

S. B. Daniels and A. Chovnick. P element transposition in *Drosophila melanogaster*. Genetics, 133:623–636, 1993.

S. B. Daniels, K. R. Peterson, L. D. Strausbaugh, M. G. Kidwell, and A. Chovnick. Evidence for horizontal transmission of the P transposable element between *Drosophila* species. Genetics, 124(2):339–55, 1990.

A. de Vanssay, A.-L. Bougé, A. Boivin, C. Hermant, L. Teysset, V. Delmarre, C. Antoniewski, and S. Ronsseray. Paramutation in Drosophila linked to emergence of a piRNA-producing locus. Nature, 490(7418): 112–115, 2012.

C. Duc, M. Yoth, S. Jensen, N. Mouniée, V. C. Bergman, Casey M, and E. Brasset. Trapping a somatic endogenous retrovirus into a germline piRNA cluster immunizes the germline against further invasion. Genome Biology, 20(127):1–14, 2019.

W. R. Engels. P elements in *Drosophila melanogaster*. In D. E. Berg and M. M. Howe, editors, Mobile DNA, chapter 16, pages 437–484. American Society for Microbiology, Washington, D.C., 1989.

W. R. Engels, D. M. Johnson-Schlitz, W. B. Eggleston, and J. Sved. High-frequency P element loss in Drosophila is homolog dependent. Cell, 62(3):515–525, 1990.

D. J. Finnegan. Eukaryotic transposable elements and genome evolution. Trends in Genetics, 5(4):103–107, 1989.

O. Gascuel. Bionj: an improved version of the nj algorithm based on a simple model of sequence data. Molecular biology and evolution, 14(7):685–695, 1997.

G. B. Gloor, N. A. Nassif, D. M. Johnson-schlitz, C. R. Preston, and W. R. Engels. Targeted Gene Replacement in Drosophila via P Element-Induced Gap Repair. Science, 253(5):1110–1117, 1991.

C. Goriaux, E. Théron, E. Brasset, and C. Vaury. History of the discovery of a master locus producing piRNAs: The flamenco/COM locus in Drosophila melanogaster. Frontiers in Genetics, 5:1–8, 2014.

L. S. Gunawardane, K. Saito, K. M. Nishida, K. Miyoshi, Y. Kawamura, T. Nagami, H. Siomi, and M. C. Siomi. A slicer-mediated mechanism for repeat-associated siRNA 5’ end formation in Drosophila. Science, 315(5818):1587–1590, 2007.

R. J. Hijmans. geosphere: Spherical Trigonometry, 2017. URL https://CRAN.R-project.org/package=geosphere. R package version 1.5-7.

D. Houle and S. V. Nuzhdin. Mutation accumulation and the effect of copia insertions in *Drosophila melanogaster*. Genetics Research, 83(1):7–18, 2004.

A. Hua-Van, A. Le Rouzic, T. S. Boutin, J. Filée, and P. Capy. The struggle for life of the genome’s selfish architects. Biology Direct, 6:1–29, 2011.

M. Itoh and I. A. Boussy. Full-size P and KP elements predominate in wild *Drosophila melanogaster*. Genes & Genetic Systems, 77(4):259–259, 2002.

M. Itoh, N. Takeuchi, M. Yamaguchi, M. T. Yamamoto, and I. A. Boussy. Prevalence of full-size P and KP elements in North American populations of *Drosophila melanogaster*. Genetica, 131(1):21–28, 2007.

I. K. Jordan, L. V. Matyunina, and J. F. McDonald. Evidence for the recent horizontal transfer of long terminal repeat retrotransposon. Proceedings of the National Academy of Sciences, 96(22):12621–12625, 1999.

Jost. Gst and its relatives do not measure differentiation. Molecular Ecology, 17:4015–4026, 2008.

Kapun, M. G. Barrόn, F. Staubach, J. Vieira, D. J. Obbard, C. Goubert, O. Rota-Stabelli, M. Kankare, A. Haudry, R. A. W. Wiberg, et al. Genomic analysis of European *Drosophila melanogaster* populations on a dense spatial scale reveals longitudinal population structure and continent-wide selection. Biorxiv, page 313759, 2018.

G. H. Karpen and A. C. Spradling. Analysis of subtelomeric heterochromatin in the Drosophila minichromosome Dp1187 by single P element insertional mutagenesis. Genetics, 132(3):737–753, 1992.

E. S. Kelleher. Reexamining the P-Element Invasion of *Drosophila melanogaster* Through the Lens of piRNA Silencing. Genetics, 203(4):1513–1531, 2016.

E. S. Kelleher, R. B. R. Azevedo, and Y. Zheng. The Evolution of Small-RNA-Mediated Silencing of an Invading Transposable Element. Genome Biology and Evolution, 0(0):evy218, 2018.

M. G. Kidwell. Evolution of hybrid dysgenesis determinants in *Drosophila melanogaster*. Proceedings of the National Academy of Sciences, 80(6):1655–1659, 1983.

M. Kimura and G. H. Weiss. The stepping stone model of population structure and the decrease of genetic correlation with distance. Genetics, 49(4):561–576, 1964. doi: 10.1093/oxfordjournals.molbev.a025590.

R. Kofler. Dynamics of transposable element invasions with piRNA clusters. Molecular Biology and Evolution, 36(7):1457—-1472, 2019.

R. Kofler, T. Hill, V. Nolte, A. Betancourt, and C. Schlötterer. The recent invasion of natural *Drosophila simulans* populations by the P-element. PNAS, 112(21):6659–6663, 2015.

R. Kofler, K.-A. Senti, V. Nolte, R. Tobler, and C. Schlötterer. Molecular dissection of a natural transposable element invasion. Genome Research, 28(2):824–835, 2018.

J. B. Lack, C. M. Cardeno, M. W. Crepeau, W. Taylor, R. B. Corbett-Detig, K. A. Stevens, C. H. Langley, and J. E. Pool. The Drosophila genome nexus: a population genomic resource of 623 *Drosophila melanogaster* genomes, including 197 from a single ancestral range population. Genetics, 199(4):1229–41, 2015.

A. Le Thomas, A. K. Rogers, A. Webster, G. K. Marinov, S. E. Liao, E. M. Perkins, J. K. Hur, A. A. Aravin, and K. F. Tόth. Piwi induces piRNA-guided transcriptional silencing and establishment of a repressive chromatin state. Genes and Development, 27(4):390–399, 2013.

E. Lerat, C. Goubert, S. Guirao-Rico, M. Merenciano, A.-B. Dufour, C. Vieira, and J. González. Population specific dynamics and selection patterns of transposable element insertions in European natural populations. Molecular Ecology, 28(6):1506–1522, 2019.

H. Li and R. Durbin. Fast and accurate short read alignment with Burrows–Wheeler transform. Bioinformatics, 25(14):1754–1760, 2009.

H. Li, B. Handsaker, A. Wysoker, T. Fennell, J. Ruan, N. Homer, G. Marth, G. Abecasis, and R. Durbin. The Sequence Alignment/Map format and SAMtools. Bioinformatics (Oxford, England), 25(16):2078–2079, Aug. 2009.

J. Lu and A. G. Clark. Population dynamics of PIWI-RNAs (piRNAs) and their targets in Drosophila. Genome Research, 20:212–227, 2009.

H. Machado, A. O. Bergland, R. Taylor, S. Tilk, E. Behrman, K. Dyer, D. Fabian, T. Flatt, J. Gonzalez, T. Karasov, I. Kozeretska, B. Lazzaro, T. Merritt, J. Pool, K. O’Brien, S. Rajpurohit, P. Roy, S. Schaeffer, S. Serga, P. Schmidt, and D. Petrov. Broad geographic sampling reveals predictable and pervasive seasonal adaptation in Drosophila. bioRxiv, page 337543, 2018.

T. F. Mackay. Transposable elements and fitness in *Drosophila melanogaster*. Genome, 31(1):284–295, 1989.

T. F. Mackay, R. F. Lyman, and M. S. Jackson. Effects of P Element Insertions on Quantitative Traits in *Drosophila melanogaster*. Genetics, 130:315–332, 1991.

S. Majumdar and D. C. Rio. P transposable elements in *Drosophila melanogaster*. Microbiol Spectrum, pages 484–518, 2015.

C. D. Malone and G. J. Hannon. Small RNAs as Guardians of the Genome. Cell, 136(4):656–668, 2009.

C. D. Malone, J. Brennecke, M. Dus, A. Stark, W. R. McCombie, R. Sachidanandam, and G. J. Hannon. Specialized piRNA pathways act in germline and somatic tissues of the Drosophila ovary. Cell, 137(3): 522–535, 2009.

F. Mohn, G. Sienski, D. Handler, and J. Brennecke. The rhino-deadlock-cutoff complex licenses noncanonical transcription of dual-strand piRNA clusters in Drosophila. Cell, 157(6):1364–1379, 2014.

E. Paradis, J. Claude, and K. Strimmer. APE: analyses of phylogenetics and evolution in R language. Bioinformatics, 20(2):289–290, 2004.

L. Pascua and G. Periquet. Distribution of hobo transposable elements in natural populations of *Drosophila melanogaster*. Molecular Biology and Evolution, 8(3):282–296, 1991.

J. Peccoud, V. Loiseau Cordaux, and C. Gilbert. Massive horizontal transfer of transposable elements in insects. PNAS, 114(18):4721–26, 2017.

G. Periquet, M. H. Hamelin, Y. Bigot, and A. Lepissier. Geographical and historical patterns of distribution of hobo elements in drosophila melanogaster populations. Journal of Evolutionary Biology, 2(3):223–229, 1989.

J. E. Pool. The Mosaic Ancestry of the Drosophila Genetic Reference Panel and the *D. melanogaster* Reference Genome Reveals a Network of Epistatic Fitness Interactions. Molecular biology and evolution, page msv194, 2015.

C. Post, J. P. Clark, Y. A. Sytnikova, G.-W. Chirn, and N. C. Lau. The capacity of target silencing by Drosophila PIWI and piRNAs. RNA (New York, N.Y.), 20(12):1977–86, 2014.

H. Quesneville, C. M. Bergman, O. Andrieu, D. Autard, D. Nouaud, M. Ashburner, and D. Anxolabehere. Combined evidence annotation of transposable elements in genome sequences. PLoS computational biology, 1(2):166–175, 2005.

R Core Team. R: A Language and Environment for Statistical Computing. R Foundation for Statistical Computing, Vienna, Austria, 2014. URL http://www.R-project.org/.

S. Schaack, C. Gilbert, and C. Feschotte. Promiscuous DNA: horizontal transfer of transposable elements and why it matters for eukaryotic evolution. Trends in ecology & evolution, 25(9):537–46, 2010.

G. Sienski, D. Dönertas, and J. Brennecke. Transcriptional silencing of transposons by Piwi and maelstrom and its impact on chromatin state and gene expression. Cell, 151(5):964–980, 2012.

K. Stadler. cultevo: Tools, Measures and Statistical Tests for Cultural Evolution, 2018. URL https://kevinstadler.github.io/cultevo/. R package version 1.0.2.

A. Telenti, L. C. Pierce, W. H. Biggs, J. Di Iulio, E. H. Wong, M. M. Fabani, E. F. Kirkness, A. Moustafa, N. Shah, C. Xie, et al. Deep sequencing of 10,000 human genomes. Proceedings of the National Academy of Sciences, 113(42):11901–11906, 2016.

L. Weilguny and R. Kofler. Deviate: Assembly-free analysis and visualization of mobile genetic element composition. Molecular ecology resources, 2019.

T. Wicker, F. Sabot, A. Hua-Van, J. L. Bennetzen, P. Capy, B. Chalhoub, A. Flavell, P. Leroy, M. Morgante, O. Panaud, et al. A unified classification system for eukaryotic transposable elements. Nature Reviews Genetics, 8(12):973–982, 2007.

S. Yamanaka, M. C. Siomi, and H. Siomi. piRNA clusters and open chromatin structure. Mobile DNA, 5 (1):22, 2014.

B. Y. K. Yukuhiro, K. Harada, and T. Mukai. Viability mutations induced by the P elements in *Drosophila melanogaster*. Jpn. J. Genet., 60:531–537, 1985.

V. Zanni, A. Eymery, M. Coiffet, M. Zytnicki, and I. Luyten. Distribution, evolution, and diversity of retrotransposons at the flamenco locus reflect the regulatory properties of piRNA clusters. Proceedings of the National Academy of Sciences, 110(49):19842–19847, 2013.

S. Zhang and E. S. Kelleher. piRNA-mediated silencing of an invading transposable element evolves rapidly through abundant beneficial de novo mutations. bioRxiv, page 611350, 2019.

