## supplement for "Reconstructing the invasion route of DNA transposons using extant population samples"

#### Supplementary figures

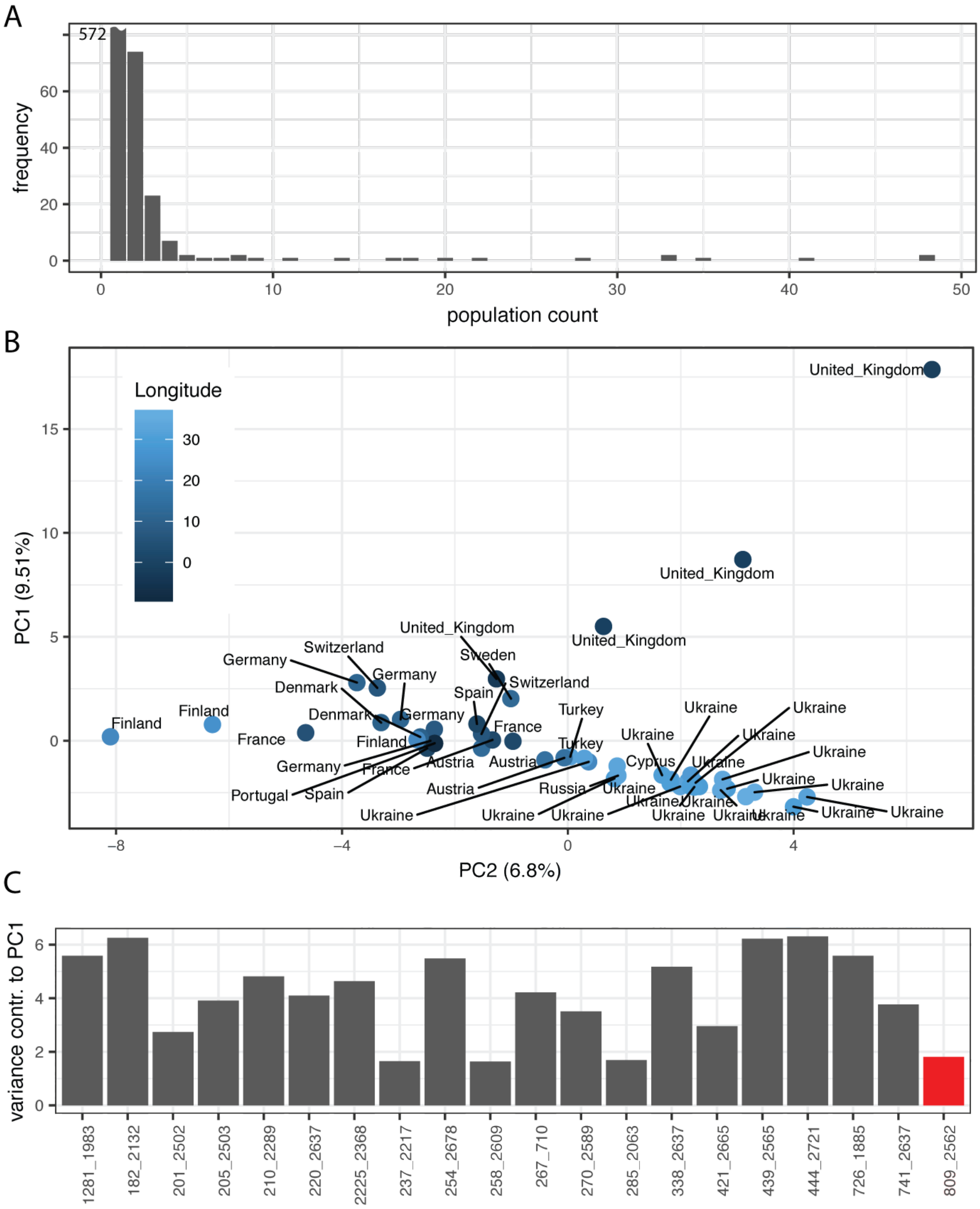


Figure S1: IDs of the P-element allow to reproduce the longitudinal cline found in *D. melanogaster* populations from Europe [Kapun et al., 2018]. A) Distribution of IDs in the investigated populations. Most IDs (572) are only found in a single population while two are found in all 48 populations. B) Scaled PCA based on the frequency of IDs found in the populations. C) Variance contribution of the most important IDs to PC1. Labels on the x-achsis indicate the start and the end position of IDs. The KP-element is shown red.

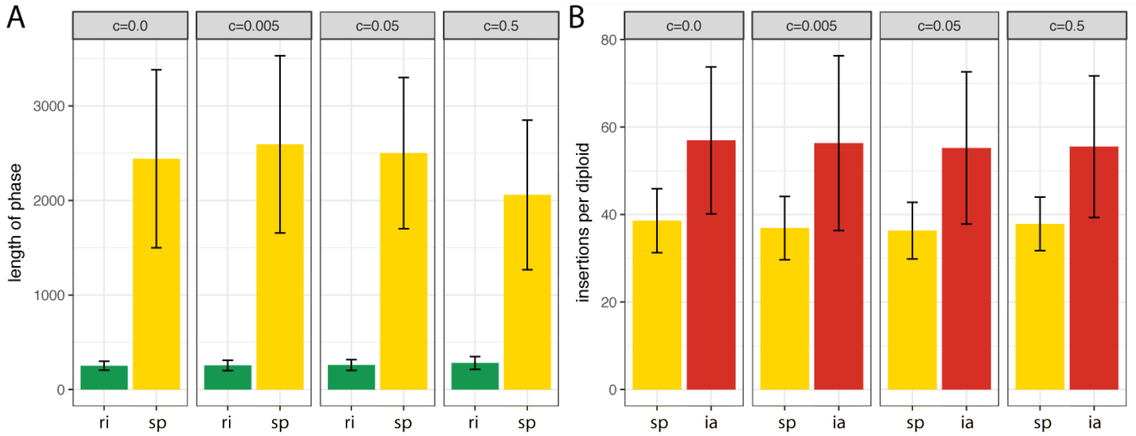


Figure S2: The probability that transposition of a FL element yields an ID (i.e. the conversion rate $c$) has little influence on the invasion dynamics. A) The conversion rate $c$ has little influence on the length of a phase (Kruskal-Wallis rank sum test; $p_{ri} = 0.11$, $p_{sp} = 0.020$). B) The conversion rate $c$ has little influence on the TE abundance at the beginning of a phase (Kruskal-Wallis rank sum test; $p_{sp} = 0.35$, $p_{ia} = 0.97$). ri rapid invasion phase, sp shotgun phase, ia inactive phase

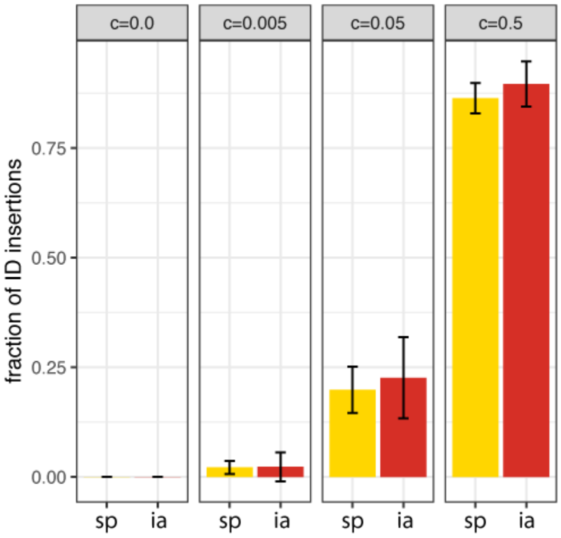


Figure S3: Once an invasion is silenced the fraction of IDs remains fairly stable within populations. We measured the fraction of IDs at the onset of the shotgun phase (sp; silencing of TEs by segregating cluster insertions) and the inactive phase (ia; silencing of TEs by fixed cluster insertions). The difference in the fraction of IDs is small and mostly not significant (Wilcoxon rank sum test; $p_{c:0.005} = 0.15$, $p_{c:0.05} = 0.22$, $p_{c:0.5} = 0.0011$). sp shotgun phase, ia inactive phase

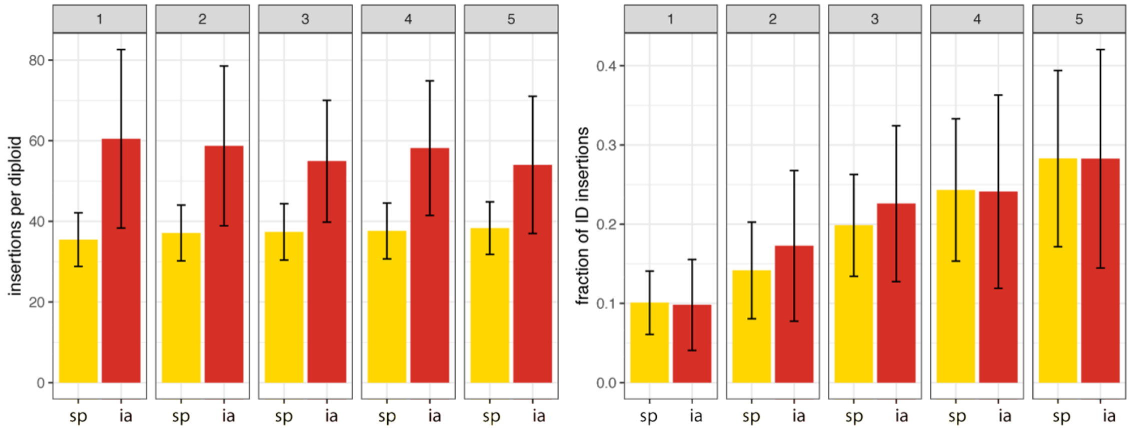


Figure S4: Although the TE abundance does not change in successively invaded populations the fraction of TEs with an ID increases in successively invaded population. Top panel indicates the order in which the populations were invaded. A) The TE abundance, measured at the beginning of two phases, remains stable in successively invaded populations (Kruskal-Wallis rank sum test; $p_{sp} = 0.19$, $p_{ia} = 0.58$) B) The fraction of IDs, measured at the beginning of two phases, increases in successively invaded populations (Kruskal-Wallis rank sum test; $p_{sp} < 2.2e-16$, $p_{ia} = 1.5e-13$). sp shotgun phase, ia inactive phase

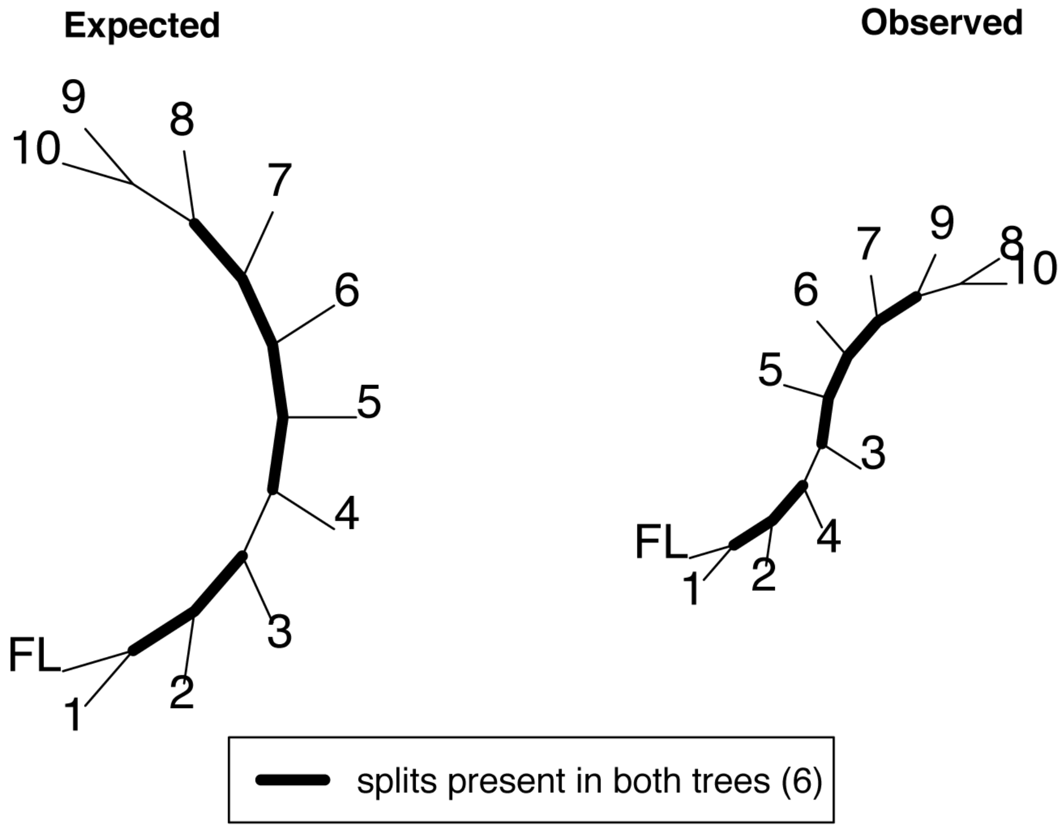


Figure S5: The number of shared splits can be used to assess the similarity between an expected and an observed tree.

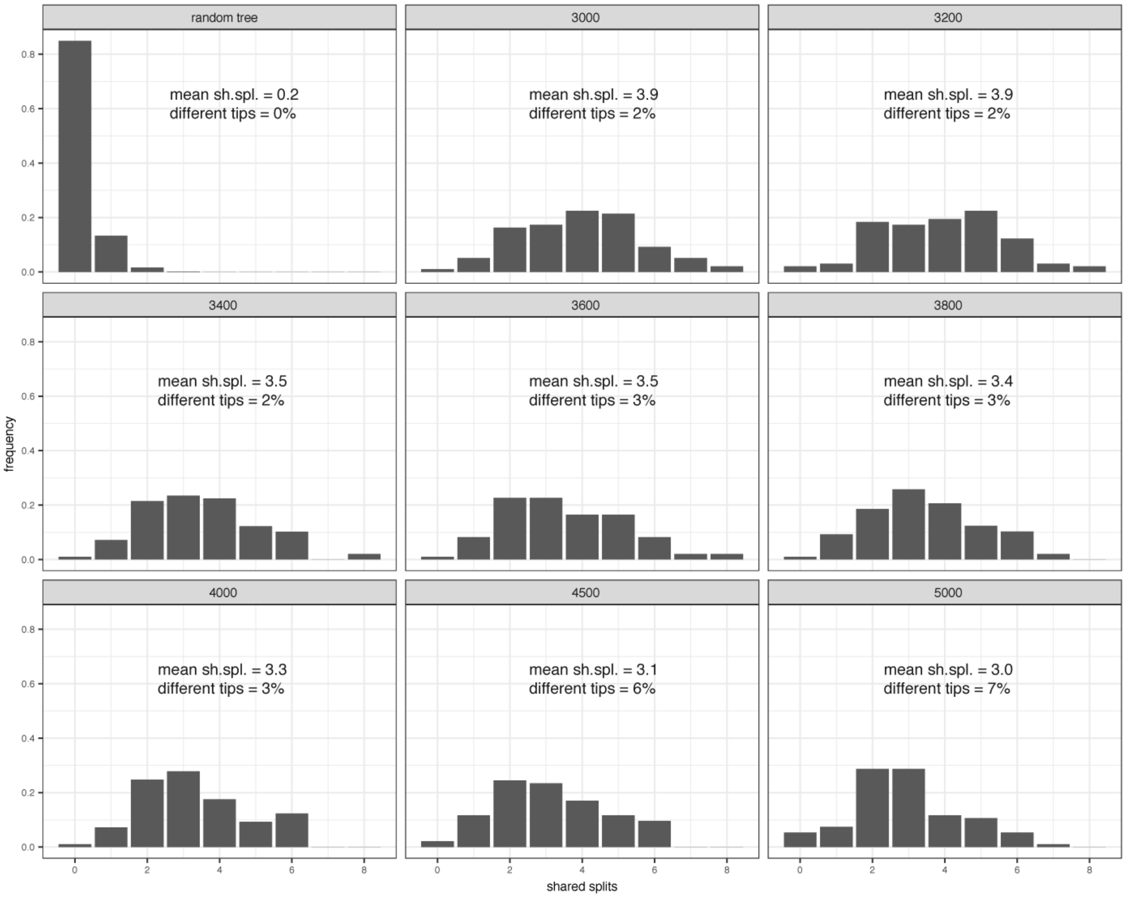


Figure S6: Influence of the sampling time (in generations; top panels) on the accuracy of the inferred invasion route. The accuracy is measured in shared splits (sh.spl.), where the maximal possible number of shared splits is 8. By generation 3000 all 10 simulated populations were invaded by the TE (it is not feasible to infer invasion routes prior to the invasion of all samples). We performed 100 simulations for each scenario, except for the random trees, where 100,000 simulations were performed. The fraction of trees having different numbers of tips than the expected one is shown in the figure.

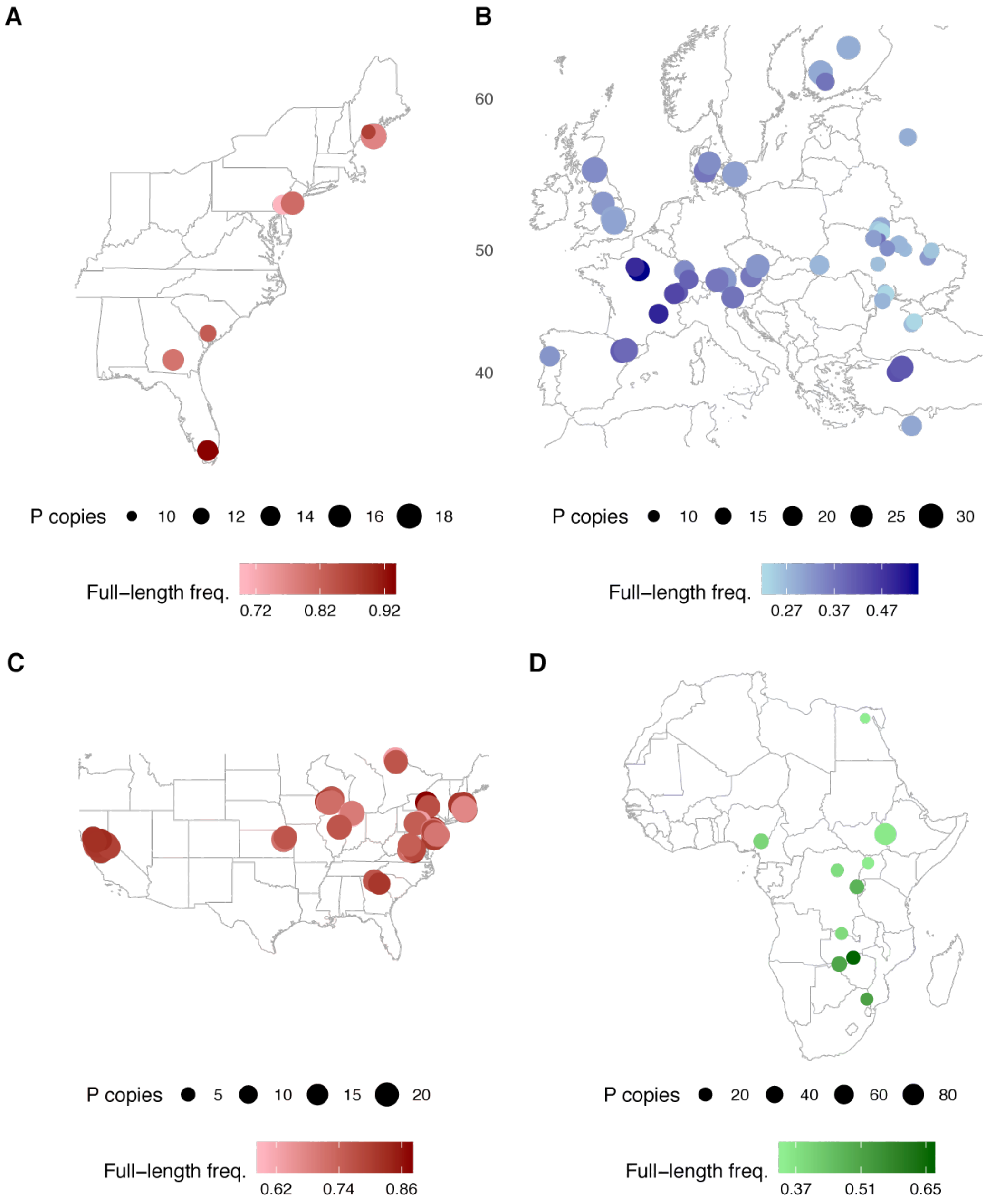


Figure S7: P-element abundance and abundance of FL elements in worldwide populations of *D. melanogaster*. Data are show for four different data sets: A) East Coast of North America [Bergland et al., 2014] B) Europe [Kapun et al., 2018] C) North America [Machado et al., 2018] and D) Africa [Lack et al., 2015]

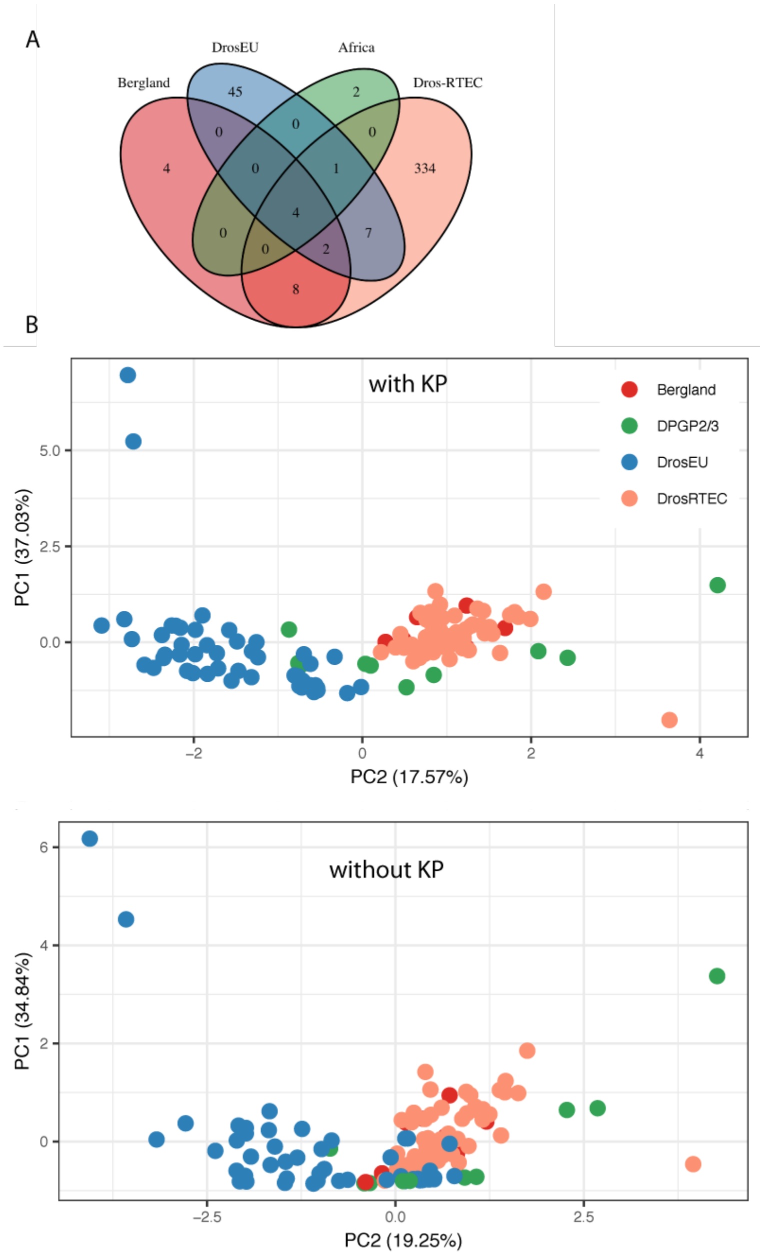


Figure S8: Overview of P-element IDs in worldwide populations of *D. melanogaster*. A) Prevalence of IDs in the four investigated data sets. Only IDs occurring in at least two populations were considered. Although most IDs are solely found in single data set, four are found in all four data sets. B) Scaled PCA based on the frequency of the six, worldwide most abundant, IDs. Note that most African samples (DPGP2/3) cluster between European (DrosEU) and North American samples (Bergland and Dros-RTEC).

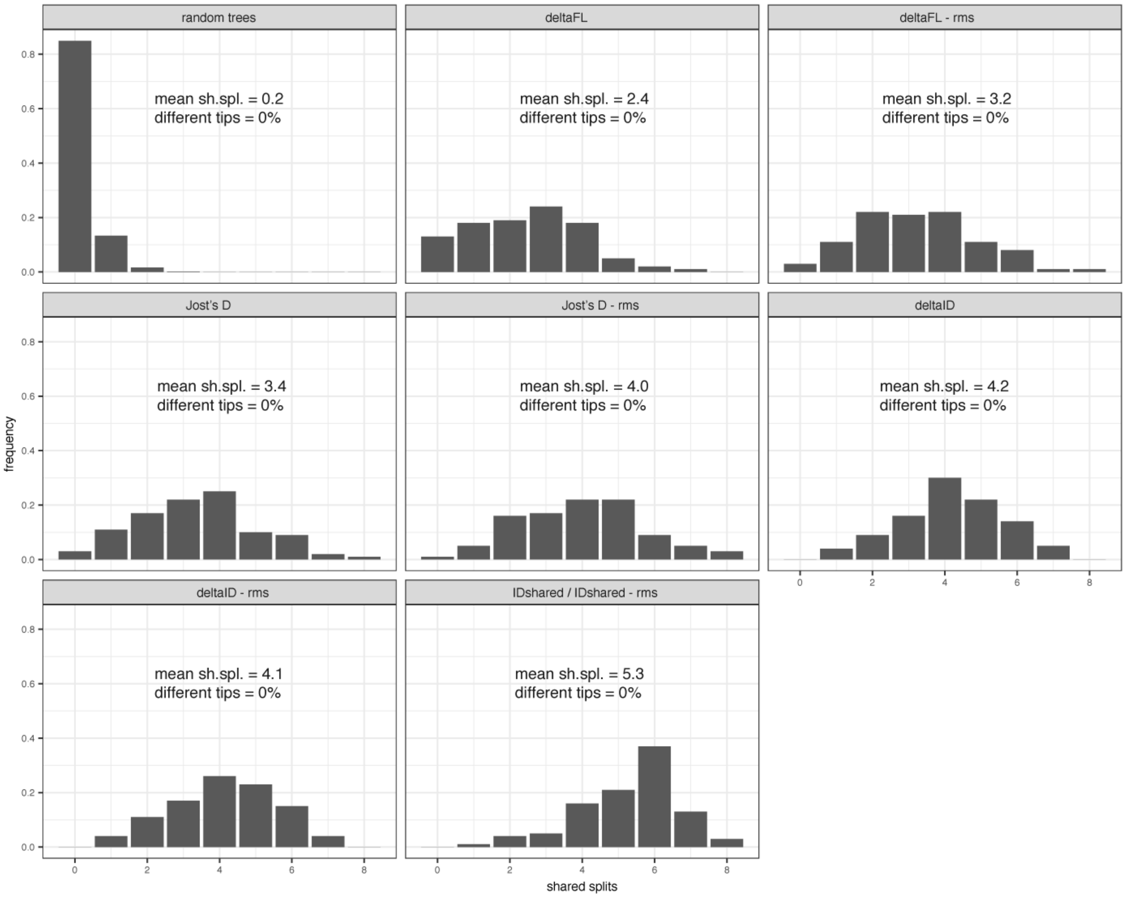


Figure S9: Influence of the test statistic on the accuracy of the inferred invasion route. The accuracy is measured in shared splits (sh.spl.), where the maximal possible number of shared splits is 8. Each test statistic was evaluated with 100 simulations and 100,000 simulations were performed for the random trees. The fraction of trees having different numbers of tips than the expected one is shown in the figure. deltaFL, difference in the fraction of FL insertions between two populations; deltaID, sum of ID frequency differences between two populations; IDshared, inverse of the number of IDs shared between two populations; rms, remove IDs solely occurring in a single population

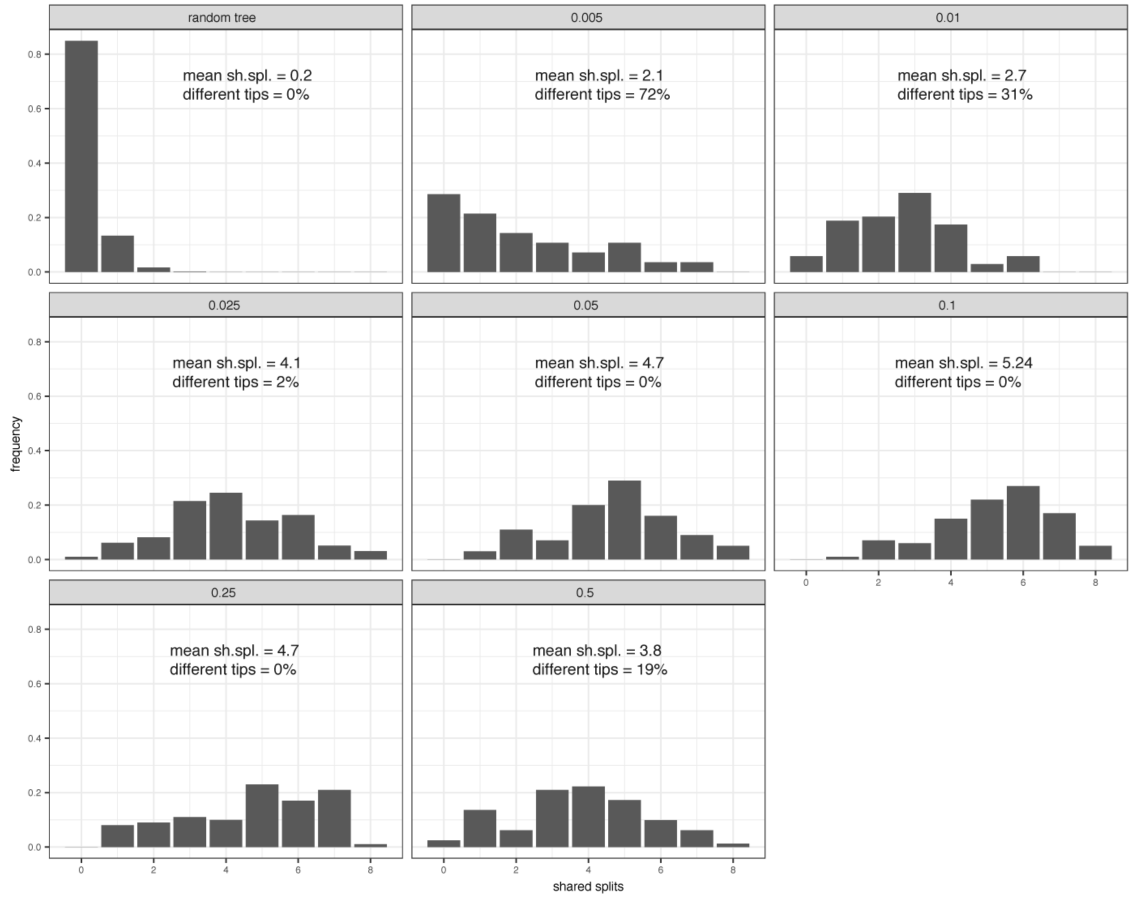


Figure S10: Influence of the conversion rate ($c$; top panel) on the accuracy of the inferred invasion route. The accuracy is measured in shared splits (sh.spl.), where the maximal possible number of shared splits is 8. We performed 100 simulations for each scenario, except for the random trees, where 100,000 simulations were performed. The fraction of trees having different numbers of tips than the expected one is shown in the figure.

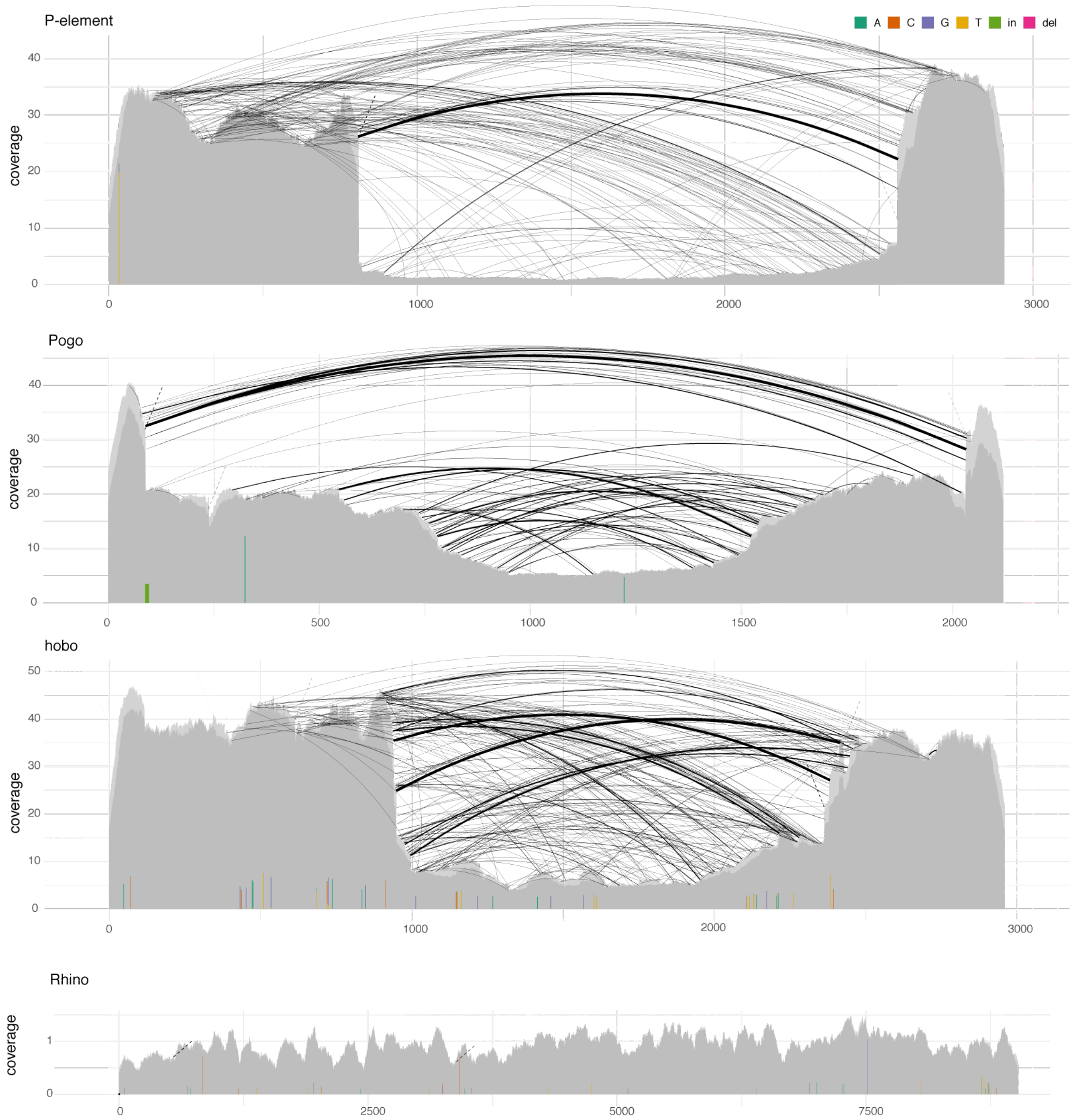


Figure S11: ID fingerprints of the P-element, Pogo and hobo. As reference the ID fingerprint of the single copy gene Rhino is shown. Data are from a natural *D. melanogaster* population sampled 2014 in Austria (Seeboden) [Kapun et al., 2018]

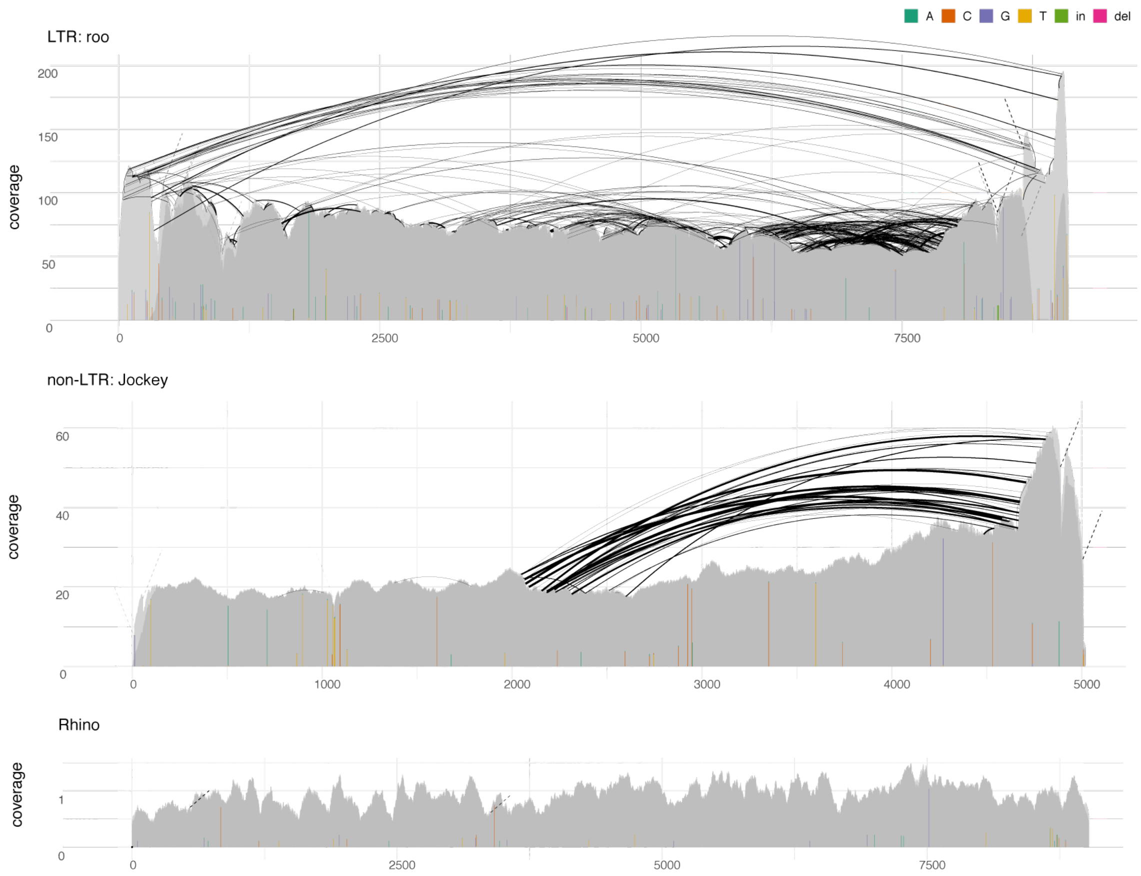


Figure S12: ID fingerprints of roo (LTR) and Jockey (non-LTR). As reference the fingerprint of the single copy gene Rhino is shown. Data are from a natural *D. melanogaster* population sampled 2014 in Austria (Seeboden) [Kapun et al., 2018]

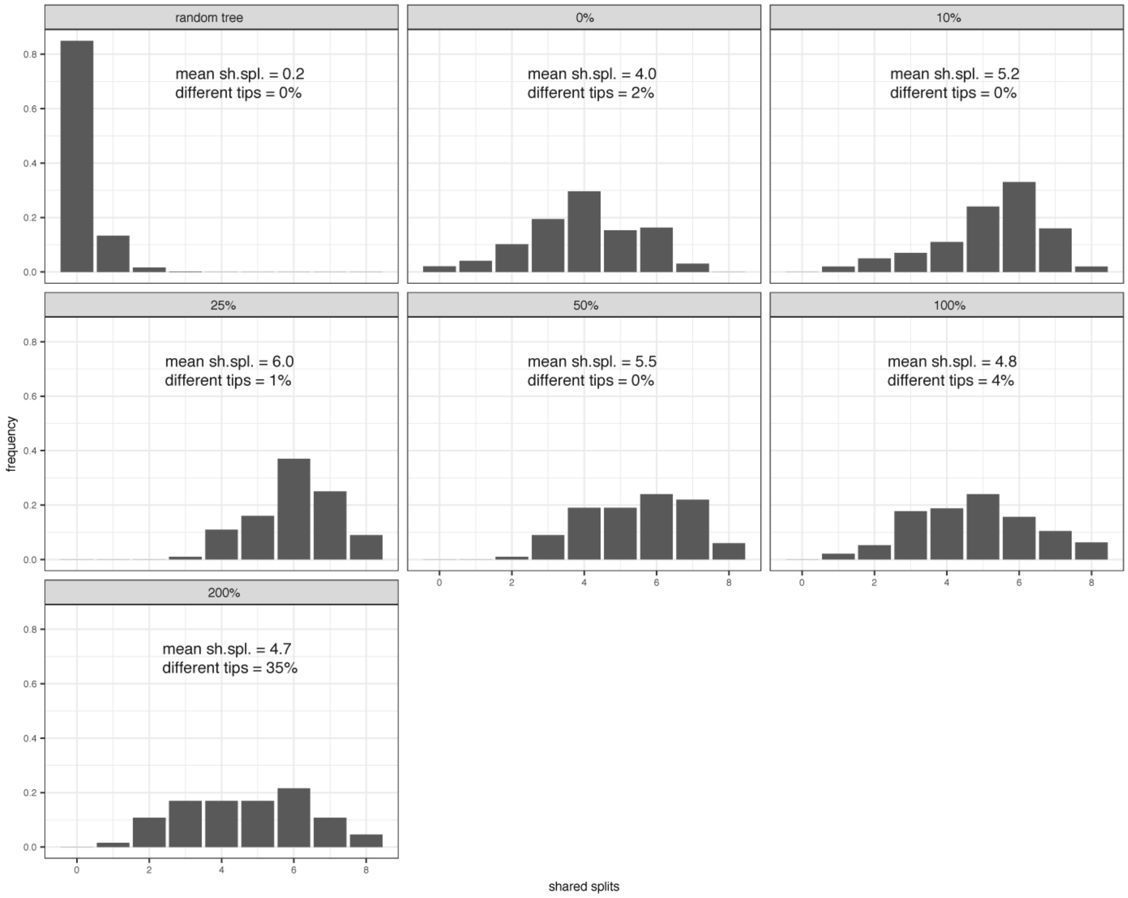


Figure S13: Influence of preferential mobilization of ID elements (in percent elevated transposition rate of ID over FL elements; top panel) on the accuracy of the inferred invasion route. The accuracy is measured in shared splits (sh.spl.), where the maximal possible number of shared splits is 8. We performed 100 simulations for each scenario, except for the random trees, where 100,000 simulations were performed. The fraction of trees having different numbers of tips than the expected one is shown in the figure.

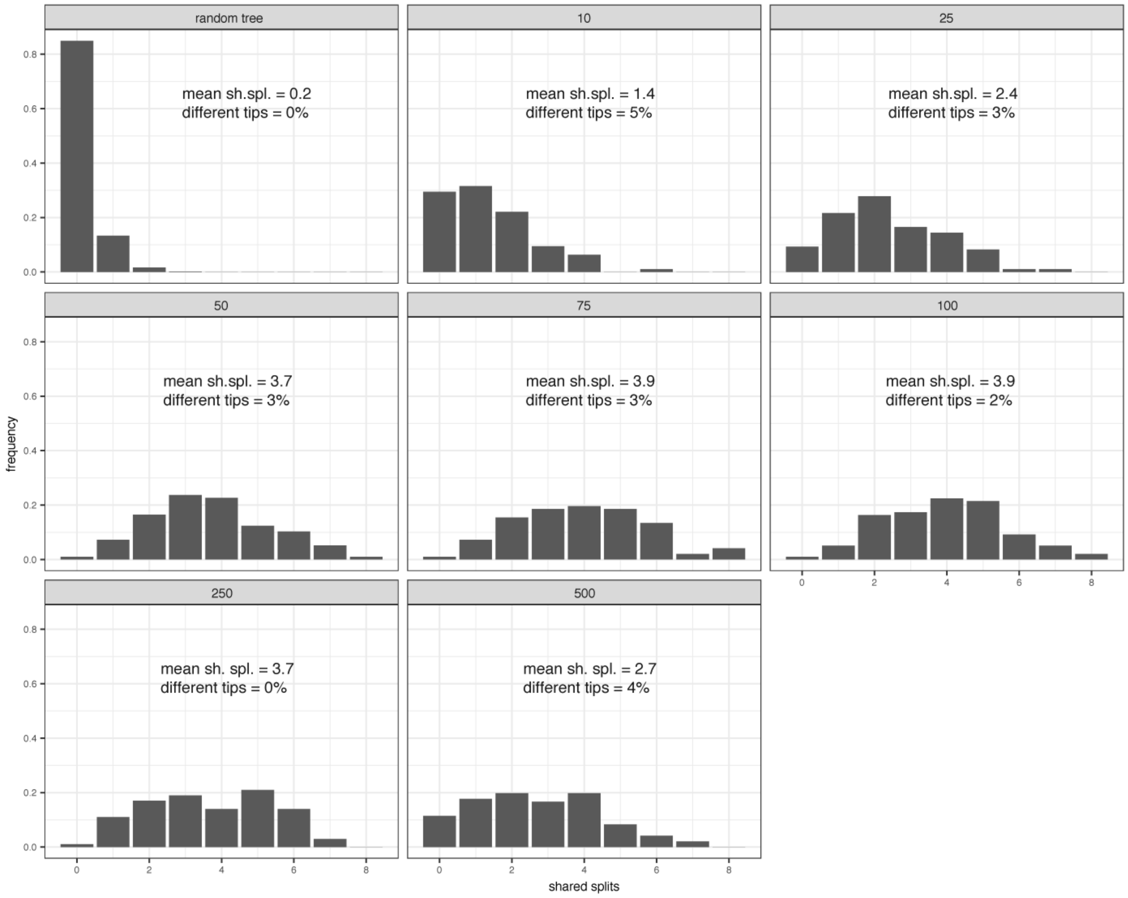


Figure S14: Influence of the number of migrants (top panel) on the accuracy of the inferred invasion route. The accuracy is measured in shared splits (sh.spl.), where the maximal possible number of shared splits is 8. We performed 100 simulations for each scenario, except for the random trees, where 100,000 simulations were performed. The fraction of trees having different numbers of tips than the expected one is shown in the figure.

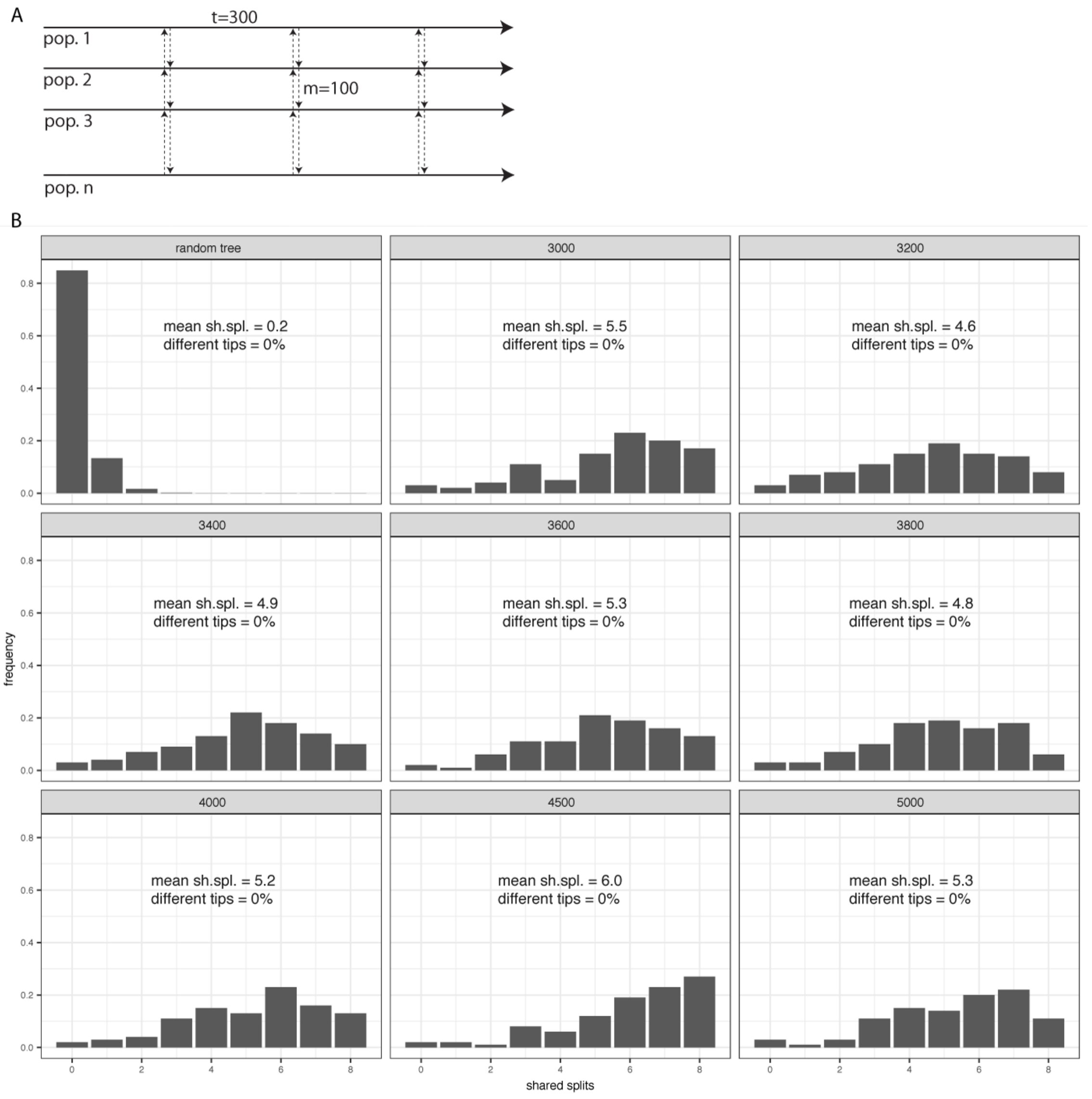


Figure S15: Influence of recurrent and bidirectional migration on the accuracy of the inferred invasion route. A) We simulated bidirectional and recurrent migration every $300^{th}$ generation between neighboring populations (with $m = 100$ migrants). B) Accuracy of the inferred invasion history in shared splits (sh.spl.) at different time points (in generations, top panel). Note that by generation 3000 all 10 simulated populations were invaded (it is not feasible to infer invasion routes prior to the invasion of all samples). The maximal possible number of shared splits is 8. We performed 100 simulations for each scenario, except for the random trees, where 100,000 simulations were performed. The fraction of trees having different numbers of tips than the expected one is shown in the figure. Compared to unidirectional-unique migration (supplementary fig. S6), bidirectional-recurrent migration increased the accuracy of our approach from 3.9 to 5.5 shared splits (at generation 3000).

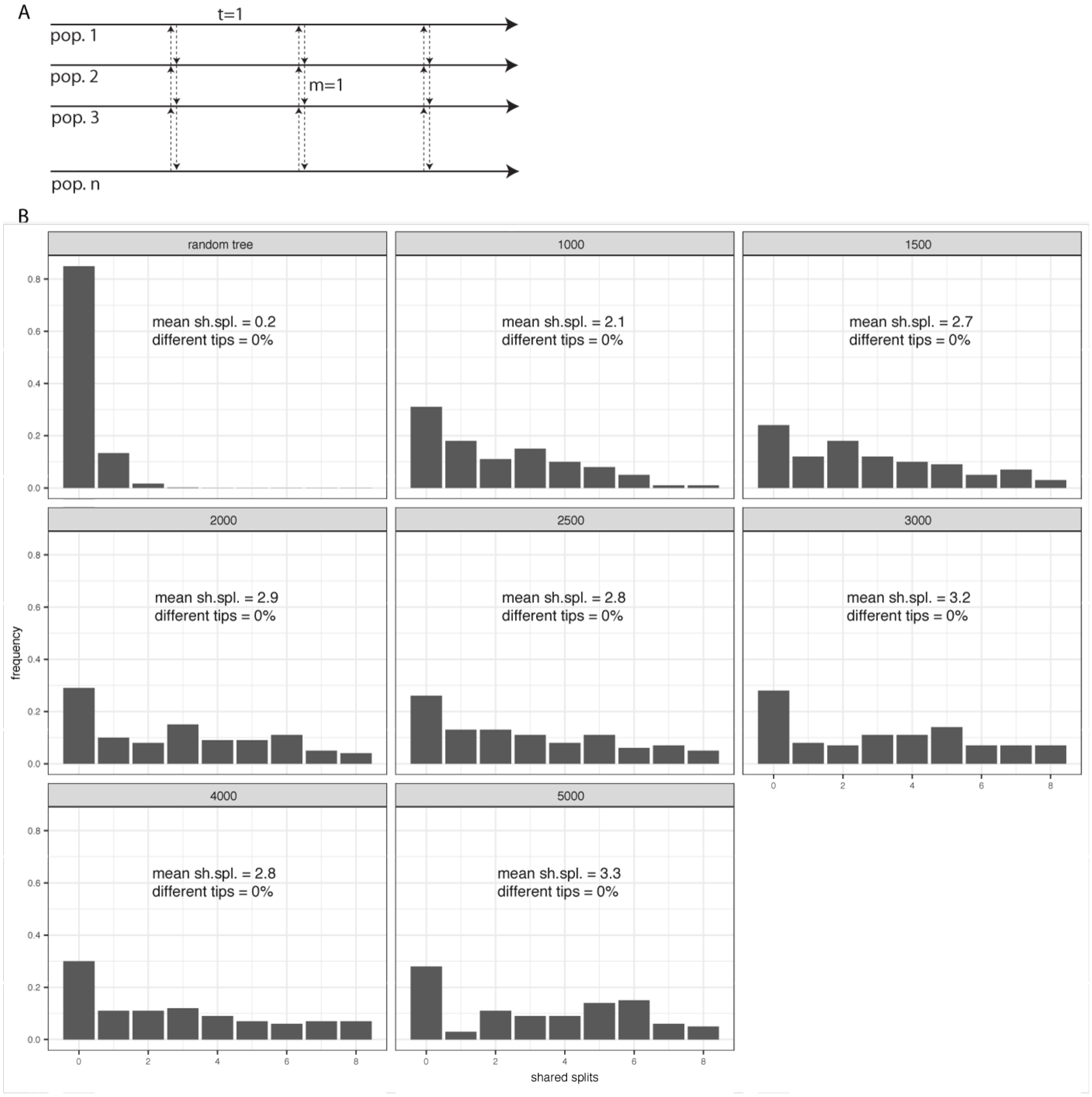


Figure S16: Influence of recurrent and bidirectional migration on the accuracy of the inferred invasion history of a TE. A) We simulated bidirectional migration of $m = 1$ migrant at each generation ($t = 1$) between neighboring populations. B) Accuracy of the inferred invasion history in shared splits (sh.spl.) at different time points (in generations, top panel). The maximal possible number of shared splits is 8. We performed 100 simulations for each scenario, except for the random trees, where 100,000 simulations were performed. The fraction of trees having different numbers of tips than the expected one is shown in the figure.

### 1 Supplementary tables

Table S1: Overview of the short read data used in this work

| publication | sample IDs |
|---|---|
| Kofler et al. [2018] | ERX2048594, ERX2048593, ERX2048592, ERX2048591, ERX2048590, ERX2048589, ERX2048588, ERX2048587, ERX2048586, ERX2048585, ERX2048584, ERX2048583, ERX2048582, ERX2048581, ERX2048580, ERX2048579, ERX2048578, ERX2048577, ERX2048576, ERX2048575, ERX2048574, ERX2048573, ERX2048572, ERX2048571, ERX2048570, ERX2048569, ERX2048568, ERX2048567, ERX2048566, ERX2048565, ERX2048564, ERX2048563, ERX2048562, ERX2048561, ERX2048560, ERX2048559 |
| Bergland et al. [2014] | SRR1525685, SRR1525694, SRR1525695, SRR1525696, SRR1525698, SRR1525699, SRR1525768, SRR1525769 |
| Kapun et al. [2018] | SRR5647729, SRR5647730, SRR5647731, SRR5647732, SRR5647733, SRR5647734, SRR5647735, SRR5647736, SRR5647737, SRR5647738, SRR5647739, SRR5647740, SRR5647741, SRR5647742, SRR5647743, SRR5647744, SRR5647745, SRR5647746, SRR5647747, SRR5647748, SRR5647749, SRR5647750, SRR5647751, SRR5647752, SRR5647753, SRR5647754, SRR5647755, SRR5647756, SRR5647757, SRR5647758, SRR5647759, SRR5647760, SRR5647761, SRR5647762, SRR5647763, SRR5647764, SRR5647765, SRR5647766, SRR5647767, SRR5647768, SRR5647769, SRR5647770, SRR5647771, SRR5647772, SRR5647773, SRR5647774, SRR5647775, SRR5647776 |
| Machado et al. [2018] | SRR3590550, SRR3590551, SRR3590554, SRR3590555, SRR3590556, SRR3590557, SRR3590558, SRR3590559, SRR3590560, SRR3590561, SRR3590562, SRR3590563, SRR3939042, SRR3939043, SRR3939044, SRR3939045, SRR3939046, SRR3939047, SRR3939048, SRR3939049, SRR3939050, SRR3939051, SRR3939052, SRR3939054, SRR3939056, SRR3939057, SRR3939058, SRR3939059, SRR3939076, SRR3939077, SRR3939078, SRR3939080, SRR3939081, SRR3939082, SRR3939083, SRR3939084, SRR3939085, SRR3939086, SRR3939087, SRR3939088, SRR3939089, SRR3939091, SRR3939092, SRR3939093, SRR3939094, SRR3939095, SRR3939096, SRR3939097, SRR3939098, SRR3939099, SRR3939100, SRR3939101, SRR3939102, SRR3939103, SRR3939104 |
| Lack et al. [2015] | SRR189040, SRR189045,SRR306633, SRR189277, SRR189279, SRR189281, SRR189389, SRR306609, SRR306611, SRR306612, SRR306614, SRR306616, SRR306618, SRR306619, SRR306621, SRR306622, SRR306623, SRR306624, SRR306629, SRR306630, SRR306631, SRR306632, SRR306634,<br>SRR202089, SRR202123, SRR203068,SRR203069, SRR203226,SRR203232,SRR203233,SRR203330,SRR203336,SRR203473,SRR203474,SRR203496,SRR203<br>SRR1686794, SRR1686796, SRR1686797, SRR1686964, SRR1688222 |

### 2 Supplementary Material and Methods

### References


Alan O. Bergland, Emily L. Behrman, Katherine R. O'Brien, Paul S. Schmidt, and Dmitri A. Petrov. Genomic Evidence of Rapid and Stable Adaptive Oscillations over Seasonal Time Scales in Drosophila. *PLoS Genetics*, 10(11):e1004775, 2014.

Martin Kapun, Maite G Barrón, Fabian Staubach, Jorge Vieira, Darren J Obbard, Clément Goubert, Omar Rota-Stabelli, Maaria Kankare, Annabelle Haudry, R Axel W Wiberg, et al. Genomic analysis of European *Drosophila melanogaster* populations on a dense spatial scale reveals longitudinal population structure and continent-wide selection. *Biorxiv*, page 313759, 2018.

Robert Kofler, Kirsten-Andre Senti, Viola Nolte, Ray Tobler, and Christian Schlötterer. Molecular dissection of a natural transposable element invasion. *Genome Research*, 28(2):824–835, 2018.

Justin B Lack, Charis M Cardeno, Marc W Crepeau, William Taylor, Russell B Corbett-Detig, Kristian A Stevens, Charles H Langley, and John E Pool. The Drosophila genome nexus: a population genomic resource of 623 *Drosophila melanogaster* genomes, including 197 from a single ancestral range population. *Genetics*, 199(4):1229–41, 2015.

Heather Machado, Alan O. Bergland, Ryan Taylor, Susanne Tilk, Emily Behrman, Kelly Dyer, Daniel Fabian, Thomas Flatt, Josefa Gonzalez, Talia Karasov, Iryna Kozeretska, Brian Lazzaro, Thomas Merritt, John Pool, Katherine O'Brien, Subhash Rajpurohit, Paula Roy, Stephen Schaeffer, Svitlana Serga, Paul Schmidt, and Dmitri Petrov. Broad geographic sampling reveals predictable and pervasive seasonal adaptation in Drosophila. *bioRxiv*, page 337543, 2018.
